# Automatic whole cell organelle segmentation in volumetric electron microscopy

**DOI:** 10.1101/2020.11.14.382143

**Authors:** Larissa Heinrich, Davis Bennett, David Ackerman, Woohyun Park, John Bogovic, Nils Eckstein, Alyson Petruncio, Jody Clements, C. Shan Xu, Jan Funke, Wyatt Korff, Harald F. Hess, Jennifer Lippincott-Schwartz, Stephan Saalfeld, Aubrey V. Weigel, COSEM Project Team

**Affiliations:** Janelia Research Campus, Howard Hughes Medical Institute, Ashburn, VA 20147, USA; Institute of Neuroinformatics UZH/ETHZ, Zurich, Switzerland

## Abstract

Cells contain hundreds of different organelle and macromolecular assemblies intricately organized relative to each other to meet any cellular demands. Obtaining a complete understanding of their organization is challenging and requires nanometer-level, threedimensional reconstruction of whole cells. Even then, the immense size of datasets and large number of structures to be characterized requires generalizable, automatic methods. To meet this challenge, we developed an analysis pipeline for comprehensively reconstructing and analyzing the cellular organelles in entire cells imaged by focused ion beam scanning electron microscopy (FIB-SEM) at a near-isotropic size of 4 or 8 nm per voxel. The pipeline involved deep learning architectures trained on diverse samples for automatic reconstruction of 35 different cellular organelle classes - ranging from endoplasmic reticulum to microtubules to ribosomes - from multiple cell types.

Automatic reconstructions were used to directly quantify various previously inaccessible metrics about these structures, including their spatial interactions. We show that automatic organelle reconstructions can also be used to automatically register light and electron microscopy images for correlative studies. We created an open data and open source web repository, OpenOrganelle, to share the data, computer code, and trained models, enabling scientists everywhere to query and further reconstruct the datasets.

---

Eukaryotic cells contain a plethora of distinct membrane-bound organelles and macromolecular assemblies that drive the cell’s activities. Despite biochemical and genetic understanding of these structures, we still lack a nanometric-level map describing their 3D distributions, morphological complexities, and interactions throughout an entire cell. Obtaining this comprehensive 3D map of cellular structures and their interactions has proved difficult. Light microscopy studies have made inroads, but the results are resolution-limited and involve potential perturbations from introduction of fluorescent protein markers^1^. Full nanometric mapping of a cell should be achievable, however, through volumetric electron microscopy of entire, unperturbed cells. Modern FIB-SEM provides near-isotropic resolution of cellular structures with nanometric voxel sizes through an entire cell^2^. Significant innovations that have been made in the automatic analysis of volume EM data^3–28^(Supplementary Information, Related Works), in turn, provide a means for accessing this information. Here, we combine whole cell FIB-SEM imaging with deep learning-based segmentation to obtain a comprehensive map of the thousands of structures packed inside a single cell.

A complete whole cell FIB-SEM dataset from a HeLa cell that was high pressure frozen and imaged at 4 nm isotropic voxels is shown in Fig. 1a. To obtain ground truth annotation for training machine learning algorithms, we selected regions of interest (hereafter, blocks) for manual annotation of up to 35 different organelles and macromolecular structures based on morphological features established in the literature. Shown in the HeLa cell of Fig. 1a (top panel) are 15 of these training blocks (see also: Supplementary Video 1). Whenever possible, we distinguished between the lumen and membrane of an organelle. Some structures were easily identified. For instance, mitochondria are ovoid organelles characterized by outer and inner membranes that fold to form cristae; microtubules are cylindrical ‘tubes’, 25 nm in diameter, that do not branch. Others can be less clear, such as the endoplasmic reticulum (ER), an extensive tubular network often studded with ribosomes. However, because it always connects back to itself and ultimately connects back to the nuclear envelope (NE) it can be confidently identified in complete volumetric EM datasets. More dubious to differentiate is the endo-lysosomal system which is often defined by its protein content - information not available in these volumetric EM. As such, we split the endosomal and lysosomal networks into two classes, organelles with a light (endosomes) and dark (lysosomes) stained lumen. Peroxisomes, also ill-defined by EM alone, are included within these two classes. For a complete list of organelle class descriptions and examples see Supplementary Methods: Organelle Classification and Extended Data Fig. 1–2.

**Figure 1 -.**
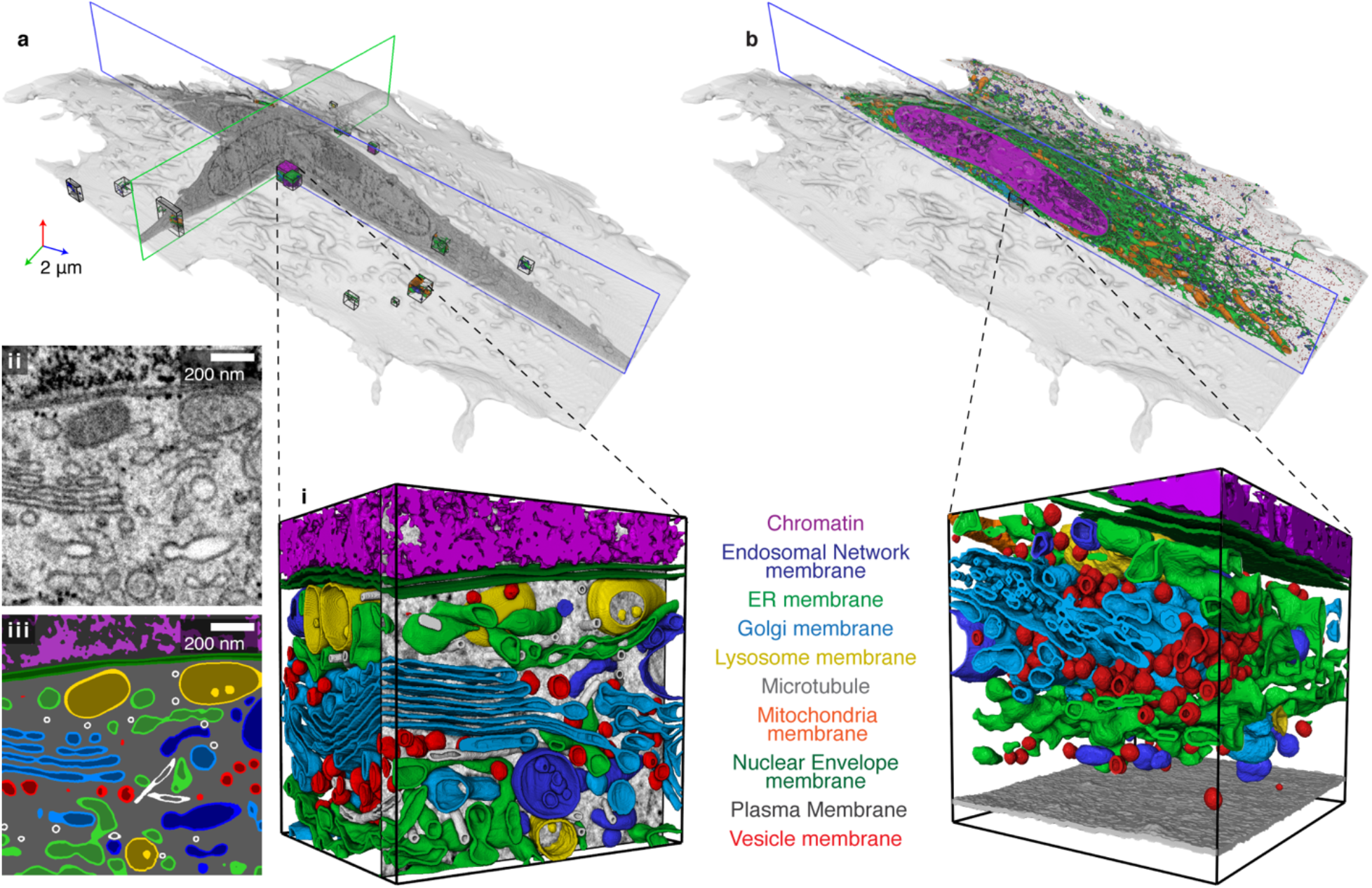
Training data and machine learning. (a) Fifteen 3D renderings of manually annotated training blocks in *jrc_hela-2.* The zoomed in subpanels demonstrate the classification of the different organelles within the volume. Shown are a single FIB-SEM slice (i) and the annotation of every voxel within the slice (ii). Alongside is a 3D rendering (iii) of a subgroup of these annotations for clarity. Bounding box is 1.2 × 1.2 × 0.95 μm. A thorough description of these classifications are reported in the Supplementary Methods. (b) Whole cell raw predictions, smoothed with σ = 18 nm and rendered with cubic interpolation, with similarly sized zoomed in region, 3 μm away from inset shown in (a). Shown are the smoothed outlines of raw organelle predictions in place of their membranes. Microtubules are left out because they were too fragmented without refinement. Bounding box is 1.2 × 1.2 × 1.2 μm.

**Figure 2 -.**
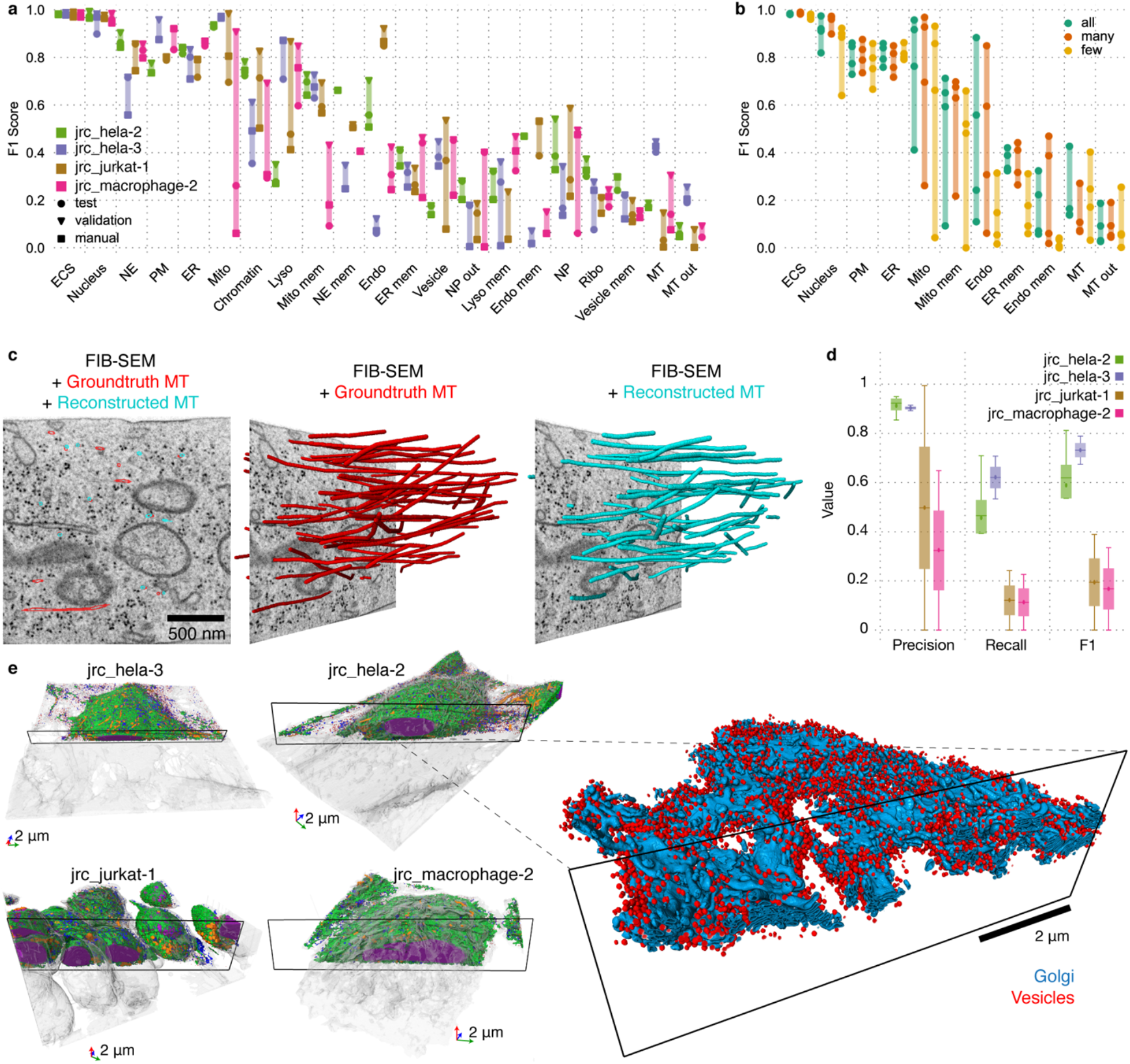
Network evaluations and refined predictions. (a) Validation and test performance measured by F1 Score on holdout blocks from four datasets. Manual validation refers to F1 Score of inferences with settings optimized manually on the whole dataset. Labels sorted by average test score. (b) Comparison of networks using different multi-class strategies. Each data point represents the F1 score (test performance) on a holdout block with the color denoting the multi-class strategy (“all”/”many”/”few”). (c) 2D FIB-SEM slice with ground truth and reconstructed microtubules after refinement in the plane, 3D renderings of the ground truth (red) and reconstructed microtubules (cyan) in a selected test block on *jrc_hela-2*. (d) Comparison of the accuracy of MT reconstruction after refinement for four different cells, measured over two densely traced 2 μm cubes for each dataset. We show reconstruction accuracy in terms of precision, recall, and F1 Score on individual edges, where an edge is correct, if the edge connects two reconstruction vertices that are matched to the same ground truth microtubule track. (e) 3D rendering of final predictions for each dataset. Classes shown are plasma membrane (gray), ER (green), mitochondria (orange), nucleus (purple), endosomal system (blue), vesicles (red), and Golgi (cyan). Inset shows Golgi and vesicles of *jrc_hela-2.*

An example of one ~1 μm^3^ area block near the cell center, with manual annotations as described above, is shown in Fig. 1a (inset i). A single FIB-SEM slice from this region, in which every voxel within the slice was annotated and classified, is shown in insets ii and iii. Segmenting the dense array of organelles in this region took one person two weeks, meaning that manual annotation of the entire cell, 2,250 times this size, would take ~60 years, underscoring the need for automatic segmentation.

The performance of deep learning-based methods depends on representative training data. Aiming for a solution of automatic segmentation that generalizes across cell types and variation in imaging parameters, we annotated up to 35 organelle classes in 28 training blocks from five datasets covering four different cell types (Extended Data Fig. 2). We further included 30 and 15 blocks that only contained extracellular space and nucleus, respectively.

The training blocks were used to train large, multi-channel 3D-U-Net architectures^29,30^ to predict signed tanh boundary distances of the binary labels for each organelle class. This was shown to be beneficial compared to using a cross-entropy loss in prior work on synaptic cleft segmentation in EM^10^. We trained all networks to minimize the sum of mean squared errors with respect to the signed tanh boundary distance of each binary label (one per output channel). Each takes on values between [-1, 1] and can be converted into binary labels by thresholding at 0. To test the effect of jointly training varying numbers of classes, one network was trained on all classes jointly (“all”), one network on the 14 classes that had positive annotations in at least 10 blocks (“many”) and 14 specialized networks on up to four closely related classes (“few”) (Supplementary Table 1). The specialized networks for some of the classes with very few positively annotated voxels in the ground truth data (i.e. ribosomes, vesicles, and Golgi) were trained with a sampling scheme that prioritized blocks containing instances of those labels.

Using the trained networks, we predicted organelle segmentation on whole cells excluding regions that contain only resin to save computation time. Depending on the network architecture, inference speeds varied between 0.5 and 4.8 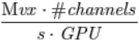 on GeForceRTX2080Ti cards. For example, on a single GPU, inference for the 14 organelles in the type “many” network on the cell shown in Fig. 1a, *jrc_hela-2* (74 gigavoxels) took approximately 12.5 hours.

We developed a manual evaluation method based on pairwise comparisons of whole-cell predictions (Supplementary Methods: Evaluations) to determine the optimal training iteration and network architecture for unseen datasets without the costly generation of validation blocks. Shown in Fig. 1b are the resulting raw predictions, smoothed with σ = 18 nm and rendered with cubic interpolation (Fig. 1b). The inset of Fig. 1b shows a region ~ 3 μm away from the inset in Fig. 1a. The quality of organelle annotations in this region, resembling those done manually in Fig. 1a(i), suggested the trained networks could successfully reconstruct organelles throughout the cell.

To quantitatively assess the performance of the machine learning method, as well as validate the manual approach, we annotated additional 4 μm^3^ holdout blocks (Extended Data Fig. 2) in four of the datasets that had annotations for training. The holdout blocks did not contain instances of all organelles, so we only report results for classes present in all four. A comparison between test and validation performances is shown in Fig. 2a. We chose the Dice coefficient (F1 Score) as our performance measure (for alternative metrics see Extended Data Fig. 3). To measure validation performance for each dataset, we optimized the aforementioned settings directly on the holdout block for that dataset. Conversely, optimizing the settings only on the holdout blocks from the other datasets allowed us to use each as a test set. Additionally, Fig. 2a shows performance scores for the settings picked via manual evaluation on the whole dataset where available (see Supplementary Table 2, for a complete comparison of manual and automatic validation see Extended Data Fig. 3). With the caveat that all holdout blocks stem from datasets that have annotations used for training, this comparison assesses the importance of hyperparameter optimization for segmentation quality. In Fig. 2b, we used the same procedure to compare test performance of the network setups (i.e. all/many/few classes). Overall, the results showed good performance scores for organelles well represented in our training data and setups including more organelles (i.e.all/many) tended to perform slightly better.

For some classes, simple refinements were applied to improve the segmentation quality. This included smoothing, size filtering, watershed segmentation and agglomeration^31,32^, and masking (see Supplementary Methods: Refinements, Extended Data Fig. 4, and SupplementaryTable 3). Evaluation of the refined segmentation performance is shown in Extended Data Fig. 5. For other classes, biological priors could be used to improve the raw predictions. In particular, for reconstructing microtubules, shown in Fig. 2c, we used the method by Eckstein et al.^28^ (Supplementary Methods: Refinements, Extended Data Fig. 6, and Supplementary Table 4). Note the close correlation between ground-truth-based (red) versus automatically reconstructed microtubules (cyan) (Fig. 2c, d).

The complete post-refinement predictions for each dataset is presented in Fig. 2e, showing seven organelles that were automatically segmented throughout four different cells. A small glimpse of the complexity of organelles in these whole cell renderings can be appreciated by focusing on the Golgi apparatus as shown in the inset of Fig. 2e: In contrast to the textbook presentations as a compact stack of cisternae, the Golgi is much more elaborate, consisting of a winding ribbon that extends > 5 μm throughout the cytoplasm and consists of flattened arrays of cisternae interconnected by narrow tubules. Vesicles are highly abundant around the Golgi, decorating its surface. For a more in depth look at the complexity of other organelles in these datasets, visit openorganelle.janelia.org.

From these whole cell datasets, we now have the ability to extract quantitative information and give a few examples of the type of analysis this data facilitates. We calculated the instance count, volume, and surface area of each organelle within each of the four different cells (Supplementary Table 5). Using the holdout blocks, we confirmed that these estimates agree with those extracted from ground truth (Extended Data Fig. 7). Fig. 3a plots the volume proportion occupied by 8 different organelle classes throughout an entire cell, comparing two HeLa cells, a Jurkat cell and a macrophage cell. Of note, the jrc_jurkat-1 cell, an immortalized T lymphocyte, had a higher fraction of its total cell volume occupied by the nucleus (~40%) compared to the other cell types, likely reflecting its origin as a leukaemic T-cell poised for proliferation. The jrc_macrophage-2 cell, a cell type involved in detection, phagocytosis, and destruction of bacteria and other agents, had an increased volume ratio of ER, which could be relevant for its extensive membrane trafficking activity.

**Figure 3 -.**
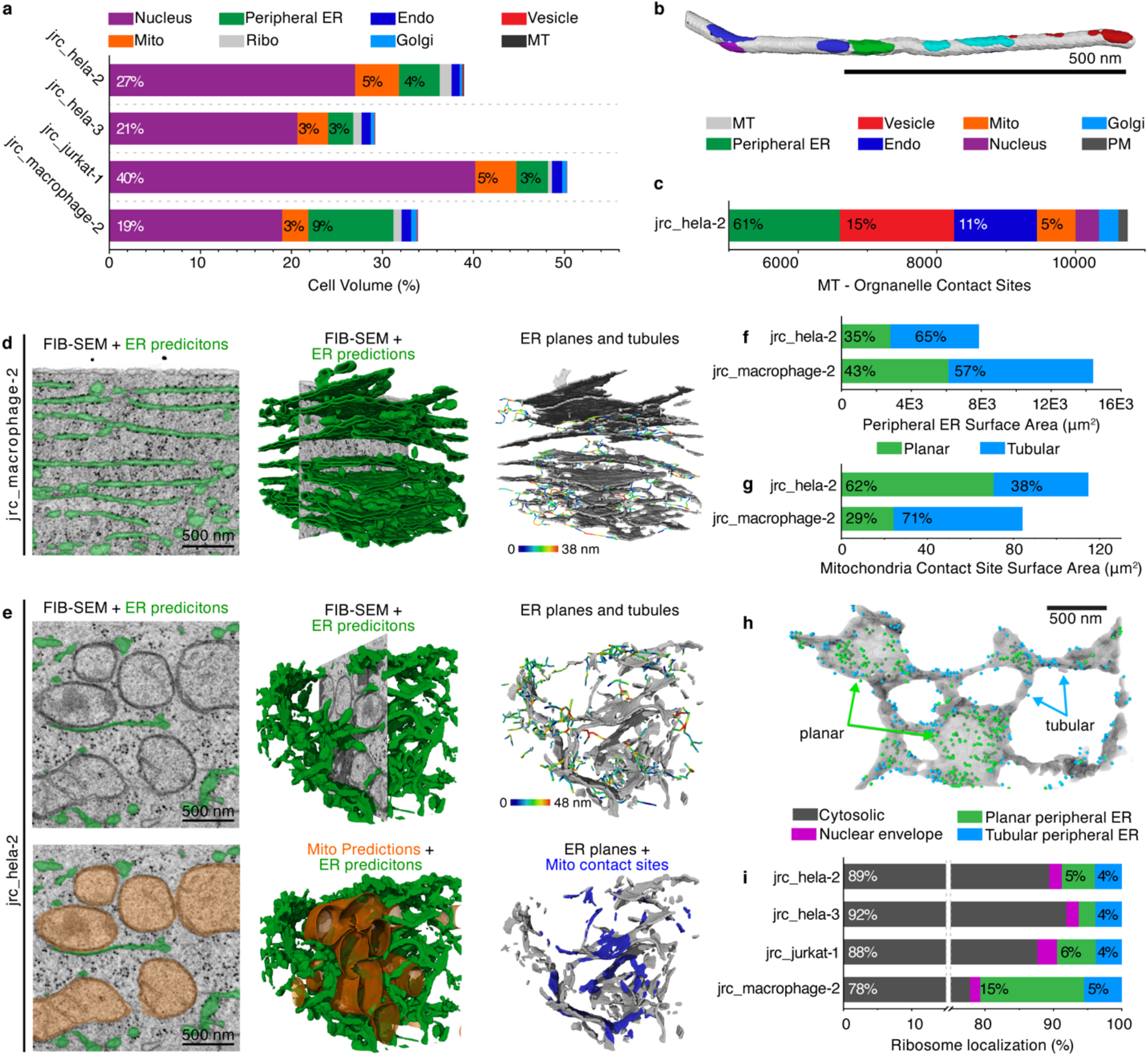
Analysis and biological insight. (a) Relative volume occupied by each predicted organelle, per cell. MT volume only shown for *jrc_hela-2*. (b) 3D rendering of reconstructed microtubule (white) and its contact sites with other organelles (colors). (c) Quantification of contact sites between microtubules and organelles in *jrc_hela-2.* (d) ER predictions in *jrc_macrophage-2.* Left panel is a 2D FIB-SEM slice with overlaid ER predictions in green, the middle panel shows a 3D rendering of the ER predictions, and for clarity the right panel shows the ER medial surface partitioned into planar and tubular structures and corresponding tubule thicknesses. (e) ER predictions in *jrc_hela-2.* Left panel is a 2D FIB-SEM slice with overlaid ER predictions in green and mitochondria predictions in orange (bottom), the middle panel shows a 3D rendering of the ER predictions and mitochondria predictions (bottom), and the right panel shows the ER medial surface partitioned into planes and tubes along with tubule thicknesses. Also shown in the bottom right panel are the contact site regions (blue) where ER and mitochondria are within 10 nm of each other. (f) Quantification of the peripheral ER curvature and surface area compared between *jrc_hela-2* and *jrc_macrophage-2*. (g) Quantification of the peripheral ER curvature at contact sites between peripheral ER and mitochondria, for *jrc_hela-2* and *jrc_macrophage-2.* (h) 3D rendering of the peripheral ER-bound ribosomes (those within 10 nm of the ER) in *jrc_hela-2.* Ribosomes are color coded based on whether they are contacting planar ER (planar measure ≥ 0.6, green) or tubular ER (planar measure < 0.6, blue). (i) Quantification of the total ribosome distribution across all four datasets based on a contact distance of 10 nm. Ribosomes that neither contact the nucleus nor ER are considered cytosolic. Peripheral ER-bound ribosomes are further categorized based on the ER curvature where they are bound.

Microtubules play a critical role in enabling organelles to distribute throughout the cell^33^. Our ability to segment out membrane-bound organelles as well as single microtubules enabled us to investigate the question of how many different organelles could be contacting a single microtubule at any time. Such associations were calculated by quantifying organelles within 20 nm of a given microtubule, as described in Supplementary Methods: Quantifications and Extended Data Fig. 8. As shown in Fig. 3b, a single microtubule, less than one micron in length, made multiple contacts with upwards of five different types of organelles (Supplementary Video 2). A summary of all the microtubule contacts in *jrc_hela-2* is summarized in Fig. 3c. These unprecedented findings open up new questions related to how organelle distributions and motions are coordinated along a single microtubule.

The peripheral ER system is known to display a variety of topologies, both within a single cell and across different cell types^34–37^. Maintaining these structural arrangements is controlled by numerous proteins ^34,35^. To quantify ER morphology, including its relationship to other organelles, we partitioned the ER into planar and tubular regions, based on its curvature (Supplementary Methods: Quantification, Extended Data Fig. 8)^36^. In the *jrc_macrophage-2* cell, 2D planar regions of ER were abundant (Fig. 3d), with many of these regions forming stacks of helicoidal sheets^37^ interconnected by tubules. In the *jrc_hela-2* cell, the flattened regions of ER often appeared to be supported by other organelles, such as mitochondria (Fig. 3e and Supplementary Video 2). Comparing the ratio of planar and tubular regions between these two cell types (Fig. 3f), we found the *jrc_hela-2* cell had fewer 2D planar regions than the *jrc_macrophage-2* cell. In line with this qualitative assessment, a larger fraction of the planar ER regions were supported by mitochondria in the *jrc_hela-2* cell (Fig. 3e, bottom right panel and Fig. 3g). These findings underscore the complexity of ER morphology and its relationship with organelles like mitochondria in different cell types.

Given the above differences between planar and tubular ER, we next examined the relative abundance of ribosomes that were associated with these ER domains. Using a contact site distance of 10 nm, we categorized ribosomes as either ER-bound or cytosolic. ER-bound ribosomes (Fig. 3h and Supplementary Video 2) were further categorized based on whether they were bound to planar ER, tubular ER, or nuclear envelope regions (Fig. 3h-i). The percentage of ribosomes bound to tubular peripheral ER and the nuclear envelope remained consistent across the data sets for the four different cells. By contrast, the relative percentage of ribosomes bound to planar peripheral ER was 3-fold higher in the *jrc_macrophage-2* cell. As the *jrc_macrophage-2* cell also had greater volume of ER than the other three cells examined (see Fig. 2a), the results are consistent with it being more actively engaged in biosynthesis of membrane and secretory proteins than the other examined cells.

While all of our training datasets were acquired at 4 nm resolution, many FIB-SEM data are acquired at 8 nm resolution to reduce imaging time. In order to leverage that data for analysis, we trained 8 nm versions of some of our setups by adding an additional upsampling block to the 3D-U-Net architecture and randomly downsampling the raw data for training (Supplementary Methods: Machine Learning). Qualitatively, segmentations of data imaged at 8 nm were satisfying, especially considering that our training data did not contain examples from these datasets. Fig. 4a shows the segmentation of another HeLa cell, which should be fairly well represented by our training data. Notably, our training data currently contains no tissue data which makes the early experiments on a sample of a chemically fixed, mouse choroid plexus^38^ (shown in Fig. 4b) particularly interesting for a future extension of the approach to tissue data and different fixation protocols. For a quantitative assessment we reused the holdout blocks with downsampled raw data as we did for training. As shown in Fig. 4c, for most organelles, the quality of segmentation suffers only slightly or, for larger organelles such as mitochondria, improves a bit compared to full resolution input.

**Figure 4 -.**
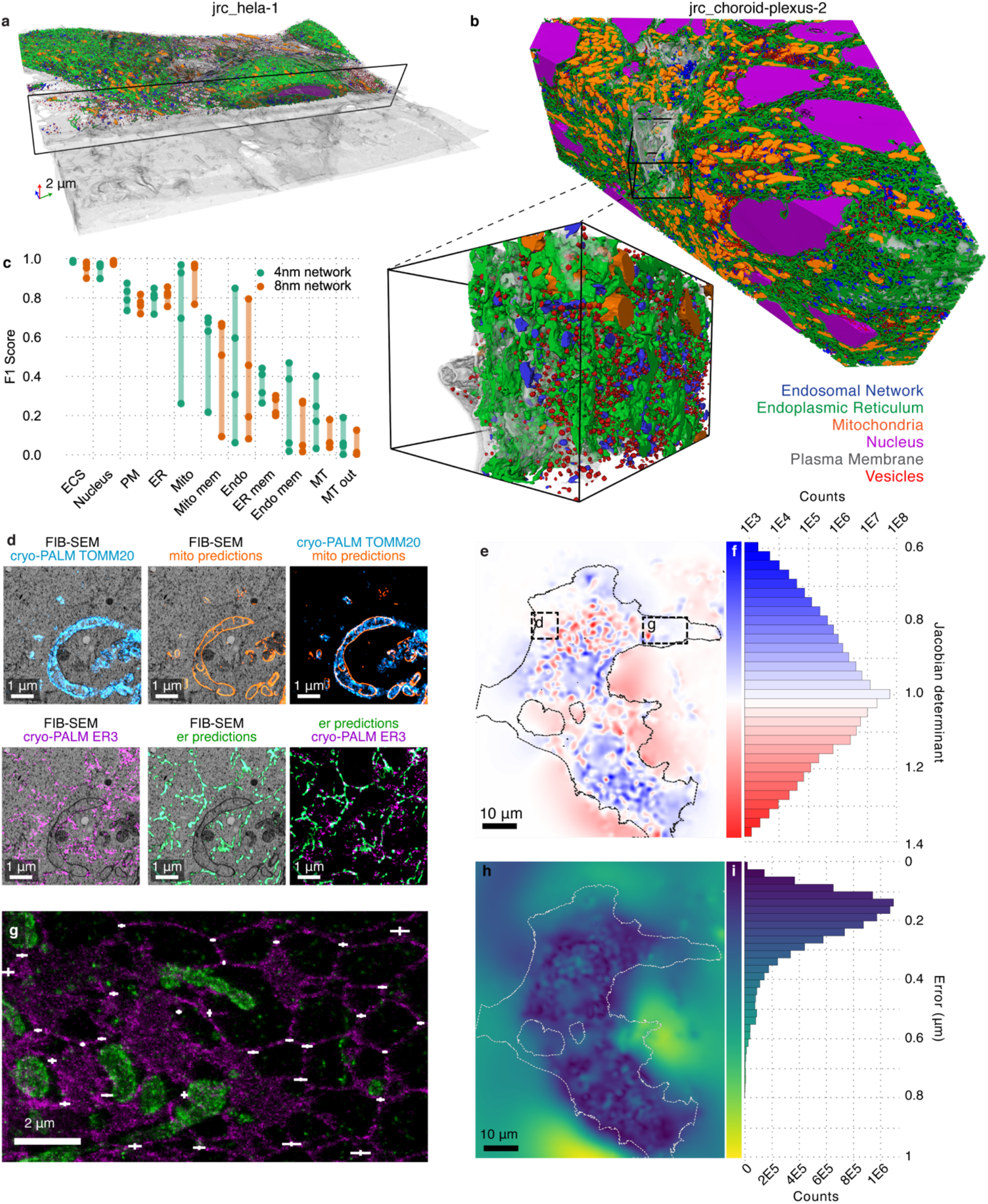
Scaling predictions and CLEM auto-registration. (a) 3D rendering of refined predictions for 8 nm dataset, *jrc_hela-1.* (b) 3D rendering of refined predictions for 8 nm dataset, jrc_choroid-plexus-2. Classes shown in (a) and (b) are plasma membrane (gray), ER (green), mitochondria (orange), nucleus (purple), endosomal system (blue), vesicles (red), and Golgi (cyan). The zoomed region is 1.2 × 1 × 0.8 μm. (c) Comparison of networks trained with 4 nm and simulated 8 nm raw data. Each data point represents the F1 score (test performance) on a holdout block. (d) Overlays of PALM images and network mitochondria / ER predictions from the region marked with a dashed box in (e). (e) A single slice of the Jacobian determinant map for the transformation registering EM to PALM for *jrc_cos7-11*. Red (blue) indicates local increase (decrease) in volume. Dotted area shows the approximate location of cells. (f) Histogram of Jacobian determinant over the whole volume. (g) Qualitative comparison of automated and manual registration for the region marked with the dashed box in (e). PALM images show ER (magenta) and mitochondria (green). White glyphs show human-human error (vertical) and human-automatic error (horizontal). (h) Error map showing differences for automatic registrations using PALM or SIM as the target image. Dotted area shows the approximate location of cells. (i) Histogram of PALM vs SIM errors over the area where a cell is present (white dotted line in (h)).

Fluorescent light microscopy (LM) in combination with highly specific molecular markers provides complementary information to high resolution non-specific EM. To associate correlative LM and EM (CLEM) acquisitions of the same sample, the acquisitions need to be registered. This is challenging due to resolution and contrast differences between the modalities, and nonlinear transformations induced during sample preparation and can take a skilled user several days to accomplish. Using our predictions for mitochondrial membranes, we automated the registration process for CLEM datasets. We demonstrate this using a previously published COS-7 cell transiently expressing ER luminal and mitochondria membrane markers, mEmerald-ER3 and Halo/JF525-TOMM20, respectively^39^, imaged by both PALM and SIM, and FIB-SEM (described in Supplementary Methods: CLEM registration). Fig. 4d, Extended Data Fig. 9 and Supplementary Video 3 show the registration results. We report the Jacobian determinant of the transformation field to measure local distortion resulting from registration. Fig. 4e shows a slice through these measurements, the cell volume is outlined for reference. Fig. 4f shows a histogram computed over the whole volume. The narrow standard deviation of this distribution (0.05) indicates that the transformation field is smooth.

We evaluated the registration accuracy with respect to human-generated ground truth by using 31 landmark pairs manually placed by two human evaluators in the subregion marked by the box in Fig. 4e. These landmarks were placed at corresponding points in the ER light channel and ER predictions of the EM image that were not used for automatic registration. This enables us to measure errors in an unbiased way, with respect to the true underlying transformation, not only the “part” of the transformation that can be inferred from the mitochondria membrane channel. Horizontal white lines in Fig. 4g shows how automated registration differs from the ground-truth human decisions, and vertical white lines show how the human evaluators differ from each other. The automatic registration differed from human annotator by 0.11 ± 0.06 μm, while human evaluators differed from each other by 0.03 ± 0.02 μm (mean ± standard deviation). The automated registration agrees with human evaluators in areas with clear mitochondria signals. Additional details and discussion can be found in the Supplementary Methods: CLEM registration.

We compared the independent registrations of the EM to both PALM and SIM. Fig. 4h shows a map of the spatial error between the two transformations. Fig. 4i shows a histogram of spatial error over a region where cells are present, indicated by the white dotted line in Fig. 4h. Errors are small, especially near mitochondria, indicating that the registration is consistent across modalities. Altogether we show that using automatic organelle segmentations can be used to successfully register CLEM images, making it accessible to less experienced users, and reducing the time from days to an hour or less.

In conclusion, large-scale reconstruction of complete cells and tissues imaged with FIB-SEM^2^ promises to greatly expand our understanding of the structure and organization of cellular organelle interactions. Fully automatic reconstruction of cellular organelle ensembles is the only economically feasible way to exhaustively analyze these large datasets. To be readily applicable to new questions and domains, it is necessary that automatic reconstruction methods generalize robustly across cell-types, tissues, and preparation methods. Until today, state-of-the-art methods are specialists that are designed for, trained on, and applied to very specific, yet often very large datasets and a small number of cellular organelles. These methods fail when presented unseen samples and require costly re-design and re-training.

We described here a first successful method to fully automatically reconstruct a large number of cellular organelles from FIB-SEM volumes of diverse cell types. We generated carefully annotated training volumes spanning a variety of different cell types and trained large deep learning architectures to simultaneously reconstruct a large number of cellular organelles and sub-organelle structures at different input resolutions.

We demonstrated that we can extract quantitative information about the distribution, size, shape and other structural properties as well as the interaction between different cellular organelles directly from automatic reconstructions of complete cells. This comprehensive information, enabling the global comparison of various cell types at the ultra-structural level, has not been previously accessible through either sparsely labeled light microscopy or focused local reconstructions on electron microscopy data. It will enable us to establish a better integrated view of cellular function and interactions between cells in tissues. We further showed that automatic organelle reconstructions enable fully automatic registration of light and electron microscopy for correlative studies, previously an error-prone manual process.

The capability of our networks to generalize across cell-types is a large step towards a pushbutton solution to analyze whole cell FIB-SEM volumes fully automatically. However, it is important to understand that more work is necessary to achieve better performance in diverse tissue samples and cell types that are vastly different from those included in our training data, and for organelles and structure that are currently underrepresented. In the Supplementary Discussion, we analyze this in detail and outline our future plans.

We strongly believe in the power of open science. We have therefore developed the repository “OpenOrganelle” (openorganelle.janelia.org) where we make raw datasets, training data, reconstructions, open source code, and models available to the public. We invite the scientific community to join us in exploring these incredibly rich datasets. Together, we will accelerate scientific discovery in both the computational and biological domain, and gain surprising insights into unexplored phenomena in cell biology.

## Methods

### Sample preparation and acquisition

Sample preparation for the four 4 nm datasets presented here are detailed in Xu et al., 2020^1^. In short, the samples were cultured on 3-mm diameter, 50 μm thick sapphire disks, high-pressure frozen, freeze-substituted with 2% OsO_4_ 0.1% UA 3% H_2_O in acetone, and resin embedded in Eponate 1. The *jrc_hela-1* dataset followed the same sample preparation. The *jrc_choroid-plexus-2* was prepared as previously described in Coulter et al., 2018^2^; postfixed in 1.0% osmium tetroxide in 0.1M cacodylate buffer (pH 7.4), rinsed with buffer, dehydrated through a graded series of ethanol, and embedded in durcupan. High-resolution (4 nm) datasets were imaged as described in Xu et al., 2020^1^ and lower-resolution datasets were imaged as described in Xu et al., 2017^3^. Note that while the actual resolution along the cutting axis can slightly differ from the lateral resolution^1^, we treated and refer to these data as isotropic. Section series were aligned using automatically extracted landmark correspondences,^4^ manually contrast adjusted, and stored at 8 bit precision using the N5 format^5^. See the Supplementary Methods: Datasets for a more detailed list of information for each dataset.

### Training data and machine learning

Organelles were manually identified using morphological features established in the literature^6^. Because every voxel within an annotated block must be classified, ‘hollow’ organelles e.g. endoplasmic reticulum, which contains lumen bound by a membrane, are defined by ‘lumen’ (lum) and ‘membrane’ (mem), or ‘inside’ (in) and ‘outside’ (out) depending on organelle composition. Examples of each class can be found in Extended Data Fig. 1.

We manually segmented blocks at 2 nm resolution from datasets of HeLa, Jurkat, Macrophage and SUM159 cells acquired at 4 nm resolution using Amira-Avizo (ThermoFisher) and BigCat^7^. The expert annotators used various image processing filters to enhance raw data, miscellaneous selection tools for label assignment and preliminary 3D rendering as a contextual guide. After final cross-annotator quality checks each block was stored as a N5 dataset^5^ in preparation for model training. For more details on the manual segmentation process and a complete list of organelles and sub-organelle compartments, see Supplementary Methods: Training Data and Supplementary Methods: Organelle Classification, respectively.

We used 3D U-Net architectures^8,9^ with valid padding and kernel sizes of (3, 3, 3) composed of 4 levels with downsampling factors of (2, 2, 2), (3, 3, 3) and (3, 3, 3) between them. The initial feature width is 12 with a multiplication factor of 6. We did not reduce the number of feature maps after the final upsampling layer. To train networks that can process data with 8 nm isotropic resolution, we added an additional upsampling level and simulated 8 nm isotropic data by sampling from 8 random subsamplings of the original data. All networks are trained with the Adam optimizer^10^ to minimize the sum of mean squared errors between the outputs and the signed tanh distance transform^11^ of each class. Distance transforms were computed on-the-fly on larger patches and then cropped to the output size of the network. Potential block boundary effects were masked out for the loss computation. We used a variety of augmentations to virtually increase the amount of training data and improve robustness to contrast variations: random flips and rotations, elastic deformations, linear and gamma intensity augmentations.

To test the trade-off between dedicated networks for each organelle and sharing information between classes, we tested networks co-predicting different numbers of classes. For a summary of all networks refer to Supplementary Table 1. Generally, we sampled patches randomly from training blocks according to their sizes. For the dedicated ribosomes, vesicles and golgi networks for 4 nm isotropic data, we sampled with probability 0.5 from the few training blocks containing instances of those. For more details on the machine learning pipeline see Supplementary Methods: Machine Learning.

Inference was performed in parallel across multiple GPUs and restricted to a mask roughly separating sample material from resin. The predicted distances were converted from float32 covering the range of [-1, 1] to uint8 with values in [0, 255], i.e. generating binary labels by thresholding the signed tanh distance transform at 0 (predicted distance of 0 nm) translates into thresholding at a voxel value of 127.

To optimize the type of network as well as the training iteration for each label and dataset we used a manual evaluation method based on repeated pairwise comparisons of random crops from two predictions. We validated this method using holdout blocks from four datasets on which we computed validation and test F1 scores by comparing to the thresholded predictions. Further details and additional metrics and comparisons can be found in the Supplementary Methods: Evaluations, Extended Data Fig. 3 and Supplementary Table 2. Refer to Supplementary Discussion for an extended discussion on manual vs automated validation.

### Refinements and quantifications

To obtain organelle instance information from the predictions, a number of refinements are performed on the predictions.

In most cases, we first smoothed the predictions with a Gaussian filter (σ = 12 nm) before thresholding at 127 and performing connected component analysis. Next, we often size-filtered the resulting segmentations to remove small false positives based on the known organelle sizes. Additional general refinements included hole filling, masking the predictions of one organelle class with those from another, and having an expert user choose specific objects to remove or keep.

For overmerged organelles we employed the watershed-agglomeration segmentation methodology of Zlateski and Seung^12^ as implemented in waterz^13^. In this approach, watershed segmentation is performed on a smoothed version of the predictions, where all background voxels are set to 0. This resulted in overly segmented organelles. To reconstruct complete organelles, iterative agglomeration was performed based on the costs of merging adjacent segments. Here, two adjacent fragments were merged if a set percentage of their shared edge voxels had predicted distances greater than or equal to a set distance; the exact percentile and distance were chosen by an expert and varied from dataset to dataset. Using this agglomeration method, fragments were more likely to be merged if their shared edge was predicted to be deep within an organelle.

Segmentation of organelles with well-defined dimensions and morphologies, such as ribosomes and microtubules, was performed by fitting the known structures to the predictions. Ribosomes are treated as spheres with radius r = 10 nm. Microtubules are particularly hard to reconstruct because they have a comparatively small diameter of 25 nm, making them hard to detect. At the same time, they extend over several microns in length, making topologically accurate reconstruction challenging. As such, for microtubule refinement, we used the method presented by Eckstein et al.^14^, exploiting structural priors on microtubules, such as no branch constraints and locally limited curvature. We found suitable hyperparameters of the method via grid search on four densely annotated 2 μm cubes of microtubules in the *jrc_hela-2* dataset. We used these hyperparameters to reconstruct all microtubules in *jrc_hela-2, jrc_hela-3, jrc_jurkat-1* and *jrc_macrophage-2* and evaluated the reconstruction accuracy for each cell on two additional, densely traced 2 μm cubes for each dataset (Fig. 2c-d). See Extended Data Fig. 6 and Supplementary Methods: Refinements for further details, including comparison of the used method to a simple baseline method. To create the 3D representation of microtubules for downstream analysis, the microtubule tracks were expanded by 12.5 nm, followed by removal of all voxels within 6 nm of the axes as well as removal of end-caps. This created tubes with an inner radius of 6 nm and outer radius of 12.5 nm.

Once the final segmentations were obtained for all organelle classes, a number of analyses were performed. For each class, we calculated instance count, volume and surface area. In addition, we measured the instance count and size of contact sites between a variety of organelle classes. We also used topological thinning^15,16^ to produce skeletons and medial surfaces of the mitochondria and ER, respectively. From the skeletons we measured mitochondrial length and diameters. From the medial surface, we partitioned the ER into planar and tubular regions based on a planarity metric. Planarity was calculated using the Hessian matrix eigenvalues for the ER, evaluated at the medial surface at a scale maximizing this metric^17^. Partitioning was then performed on a planarity-labeled version of the ER, reconstructed from the medial surface measurements, thresholded at 0.6.

For more detail on refinements and quantifications, see Supplementary Methods: Refinements, Quantifications.

### CLEM registration

We downsampled and blurred mitochondria membrane predictions to generate a “synthetic light” image. We registered this image to light images labelling Halo/JF525-TOMM20. First, a human annotator coarsely masks corresponding cells in both images for automatic processing. We used elastix^18^ to register these images in two steps: affine-only followed by deformable. For additional details, see Supplementary Methods: CLEM Registration.

### Open data and open source

We are committed to sharing the raw and derived datasets presented in this paper with the broader scientific community, for educational purposes and further analysis and discovery. Sharing data at this scale presents a technical challenge, which we addressed through a combination of technologies: we stored datasets on the cloud using Amazon Web Services (AWS) S3 storage platform using a variety of “cloud-friendly” data formats. To make this collection of datasets viewable, we built a web platform “OpenOrganelle” (openorganelle.janelia.org) that provides a catalog of datasets and access to tools for data visualization.

### Cloud Storage

We used AWS S3 for hosting our public-facing datasets, as the AWS S3 cloud storage platform offers high scalability, availability, and extensively documented application programming interfaces (APIs) across a wide range of programming languages. Additionally, AWS offers free hosting for selected open datasets, including ours, through their open data program. See https://registry.opendata.aws/janelia-cosem/ for more details about our presence on this program.

### Data formats

We stored all of our datasets in “cloud-friendly” formats, i.e. file formats that enable web services to easily access data using standard web communication protocols like hypertext transfer protocol (HTTP). For large imaging datasets a cloud-friendly format is one that provides a web-compatible interface for making spatial queries against the data. This constraint ruled out traditional bioimaging formats like mrc, tiff and hdf5 because these formats do not expose their spatial structure to HTTP. Thus we exclusively stored volumetric image data using N5 and “Neuroglancer precomputed” file formats, both of which represent image volumes as collections of separate files, or “chunks”. When hosted on a cloud storage platform like S3, each chunk has its own URL and thus can be individually accessed over HTTP. To save storage space and speed up data transmission each chunk is compressed with a browser-compatible compression scheme (GZip for lossless compression or JPEG for lossy compression).

### OpenOrganelle

Although our data can be publicly accessed directly through the cloud storage API, this is not particularly user friendly or amenable to data exploration. Thus we created a public-facing web page (www.openorganelle.ianelia.org) that provides a description of each dataset and access to tools for viewing individual subvolumes (e.g., mitochondria predictions) within a dataset. Our site does not provide any direct tools for data visualization; instead, our site dynamically creates hyperlinks that target a publicly hosted instance of the web-based image viewer Neuroglancer^19^, which is designed to access large cloud-hosted datasets over HTTP. Our site is written in TypeScript using the React^20^ framework and is completely open source (https://github.com/ianelia-cosem/openorganelle).

### Data organization

We stored all of our large volumetric datasets as image pyramids, i.e. a given volume is stored as a collection of arrays at different levels of detail. Although this approach increased the amount of storage space needed, it is the preferred format for interactively visualizing large volumes. Creating an image pyramid involves repeatedly downsampling an image volume using increasing downsampling factors. We performed downsampling by partitioning the source data into a collection of contiguous cubes of voxels with edge length 2^N^, where N ranged from 1 to 5 and computing a reducing function over the values within each cube. For image volumes with voxel values representing a scalar quantity, such as EM volumes or unrefined predictions, that reducing function was the arithmetic mean, while for volumes representing labels and segment IDs (e.g., segmentation volumes) we reduced by computing the modal value in each cube. Most of our volumes are stored in the N5 format, in which case we stored each image pyramid as a collection of datasets in a group. The group metadata contains an enumeration of the datasets that comprise the multiresolution pyramid, as well as a specification of the spatial transformation for each dataset. A minority of our volumes (those that warranted lossy jpeg compression) are stored in the Neuroglancer precomputed format, which is specific to the Neuroglancer viewer tool and natively supports image pyramids through its own metadata specification.

### Data visualization and access

Data was made available for online viewing via Neuroglancer^19,21^, an open source WebGL data viewer. Neuroglancer provides intuitive navigation of 3D datasets and supports orthogonal slice views, image pyramids, mesh rendering and a variety of other features making it ideal for displaying the large 3D datasets produced by this project. For visualization, we provide the raw data, predictions, segmentations, contact sites, skeletons and medial surfaces discussed throughout the text. For select organelles and contact sites, we also include meshes^22^ in the Neuroglancer legacy mesh format.

We provide two plugins for Fiji^23^ to access and interact with the data we provide. N5-Viewer enables viewing these large datasets in their entirety with BigDataViewer^24^. N5-ij^25^ allows users to open datasets or subsets of datasets in Fiji to perform their own analyses.

To download datasets, we recommend using the AWS command line interface (AWS CLI). Instructions for using this tool can be found on their user guide^26^.

We have also made data analysis results available as a Neo4j graph database (downloadable from openorganelle.ianelia.org). Neo4j allows users to easily interact with and visualize the data, including organelle properties like volume and surface area and contact site properties like contact area and planarity. Neo4J also allows users to create custom queries to search through and analyze data and relationships in novel ways.

We welcome feedback about how these data are used and your biological discoveries. We hope this will inspire new insights and expect they will generate more questions than answers. Please email us at

## Data Availability

All data, software, and source code generated and analyzed during this study can be found at https://openorganelle.ianelia.org/.

## Acknowledgements

This work is part of the <u>COSEM Project Team</u> at Janelia Research Campus, Howard Hughes Medical Institute, Ashburn, VA.

During this effort, the COSEM Project Team consisted of: Riasat Ali, Rebecca Arruda, Rohit Bahtra, Davis Bennett, Destiny Nguyen, Woohyun Park, Alyson Petruncio, led by Aubrey Weigel with Steering Committee of Jan Funke, Harald Hess, Wyatt Korff, Jennifer Lippincott-Schwartz, and Stephan Saalfeld.

We thank Rohit Bahtra, a Janelia-LCR Summer Internship Program student, for his work generating masks of datasets and providing manual annotations.

We thank Arslan Aziz, a Janelia Undergraduate Scholars Program student, for his work correcting mitochondria over-merging.

We thank Gudrn Ihrke and Project Technical Resources for management and coordination and staff support.

We thank Janelia Scientific Computing Shared Resource, especially Tom Dolafi and Stuart Berg for their help generating the database and visualization tools.

We thank Constantin Pape and Juan Nunez-Iglesias for their work on the inference pipeline.

We thank Victoria Custard for administrative support.

We thank Schuyler van Engelenburg, Huxley Hoffman, Eric Betzig, David Hoffman, Christopher

Walsh, and Michael Coulter for graciously providing their data.

We thank Amazon Web Services for free hosting of our data through their open data program.

We thank Song Pang and Gleb Shtengel for their work collecting and organizing the FIB-SEM data to seed OpenOrganelle. We further thank Gleb Shtengel for his early work manually segmenting organelles, motivating the need for more automated approaches.

We thank Geoffrey Meissner for critical reading of the manuscript.

Three of the datasets used in this research were derived from a HeLa cell line. Henrietta Lacks, and the HeLa cell line that was established from her tumor cells without her knowledge or consent in 1951, have made significant contributions to scientific progress and advances in human health. We are grateful to Henrietta Lacks, now deceased, and to her surviving family members for their contributions to biomedical research.

This work was supported by Howard Hughes Medical Institute, Janelia Research Campus.

## Author contributions

C.P.T, S.S., A.V.W., W.K., J.L.-S., H.F.H., and J.F. conceptualized the project.

C. S.X. and H.F.H. provided the FIB-SEM data and pre-processing.

W.P., A.P., and A.V.W. provided manual annotations, evaluations, and proofreading.

D. B. built the data management infrastructure.

L.H., and S.S. developed machine learning algorithms; L.H. performed network training and automatic evaluations.

N.E., and J.F. developed MT modeling algorithms; N.E. performed MT modeling.

D.A., and S.S. developed refinement and analysis algorithms; D.A. analyzed data.

J.B., and S.S. developed automated CLEM registration algorithms; J.B. performed automated CLEM registration.

D.B., J.C., S.S., and A.V.W. developed the data portal, OpenOrganelle.

L.H., A.V.W., S.S., D.B., D.A., J.B., N.E., and A.P. wrote the manuscript with input from all coauthors. J.L.-S. provided critical review, commentary and revision in the writing process of the manuscript.

COSEM Project Team supervised the project with guidance from S.S., J.L.-S., H.F.H., J.F., and W.K.

## Competing interests

The authors declare no competing interests.

## Correspondence

Correspondence to Aubrey Weigel and Stephan Saafleld.

## Additional information

Supplementary Information file includes Supplementary information text, methods,

Supplementary Videos 1-3 and Supplementary Tables 1-5.

## Extended data figures

**Extended Data Fig. 1 -.**
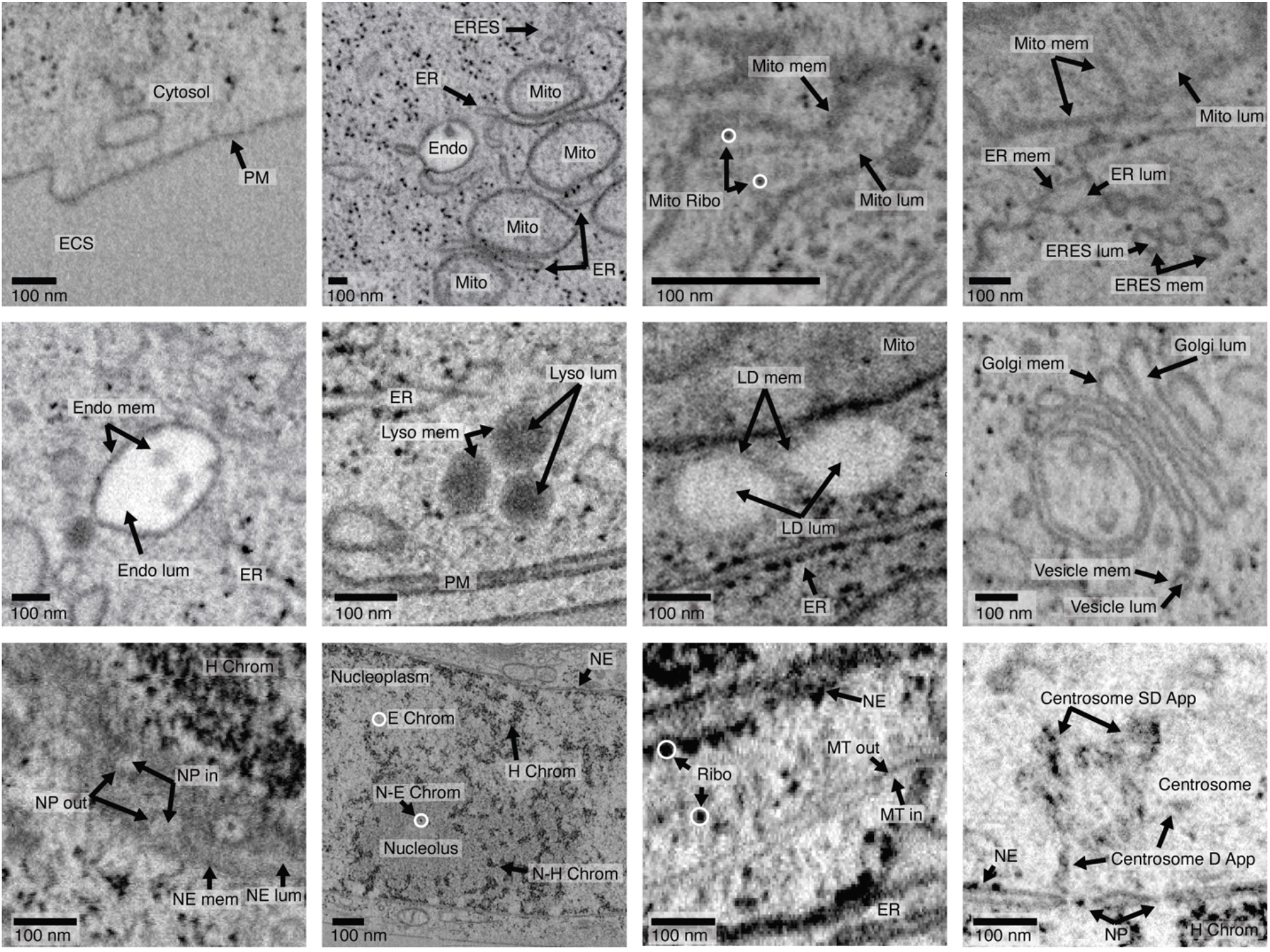
Organelle classification. Examples of each class used for training input. Organelles were manually identified using morphological features established in the literature. A description of each class can be found in the Supplementary Methods: Organelle classification.

**Extended Data Fig. 2 -.**
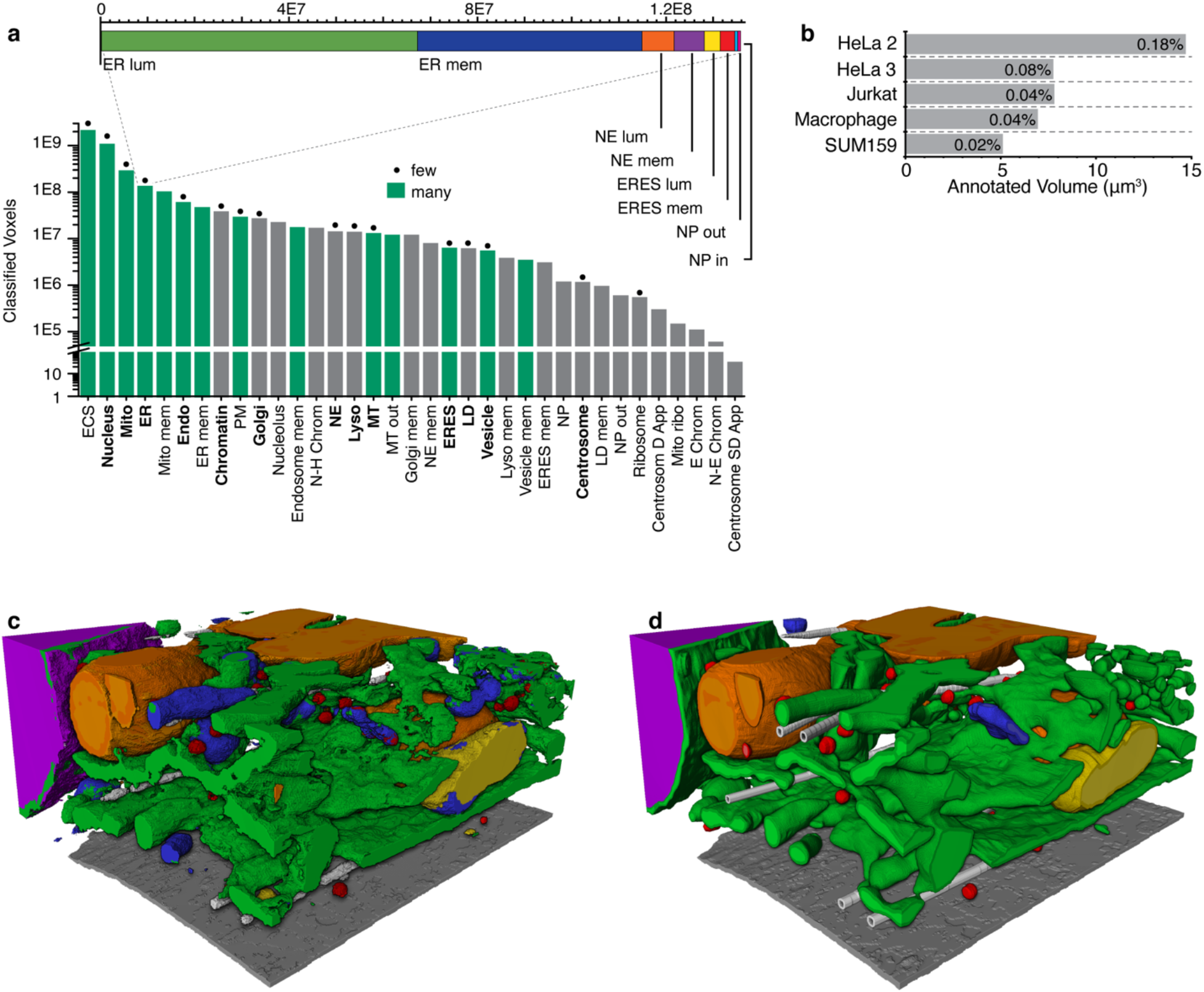
Class frequencies and holdout block. (a) 37 different classes are used to classify all intracellular structures. These classes are combined into 35, potentially overlapping semantic categories (see Supplementary Methods and Extended Data Fig. 1). Classes in bold depict these super classes. As an example, the ER object class is expanded in the subpanel. In green are classes that are predicted jointly by type “many” networks. Denoted with a dot are the super classes, with a few additional classes, used in the type “few” networks. The type “all” networks train jointly on all 35 classes. (b) Annotated volume according to datasets. Reported are the percentage of the total cell that is annotated. 3D rendering of predictions (c) and ground truth (d) in a 4 μm × 4 μm × 4 μm holdout block in *jrc_hela-3.* Shown are the nucleus (magenta), plasma membrane (gray), ER and NE (green), mitochondria (orange), vesicles (red), lysosomes (yellow), endosomes (blue), and microtubules (white).

**Extended Data Fig. 3 -.**
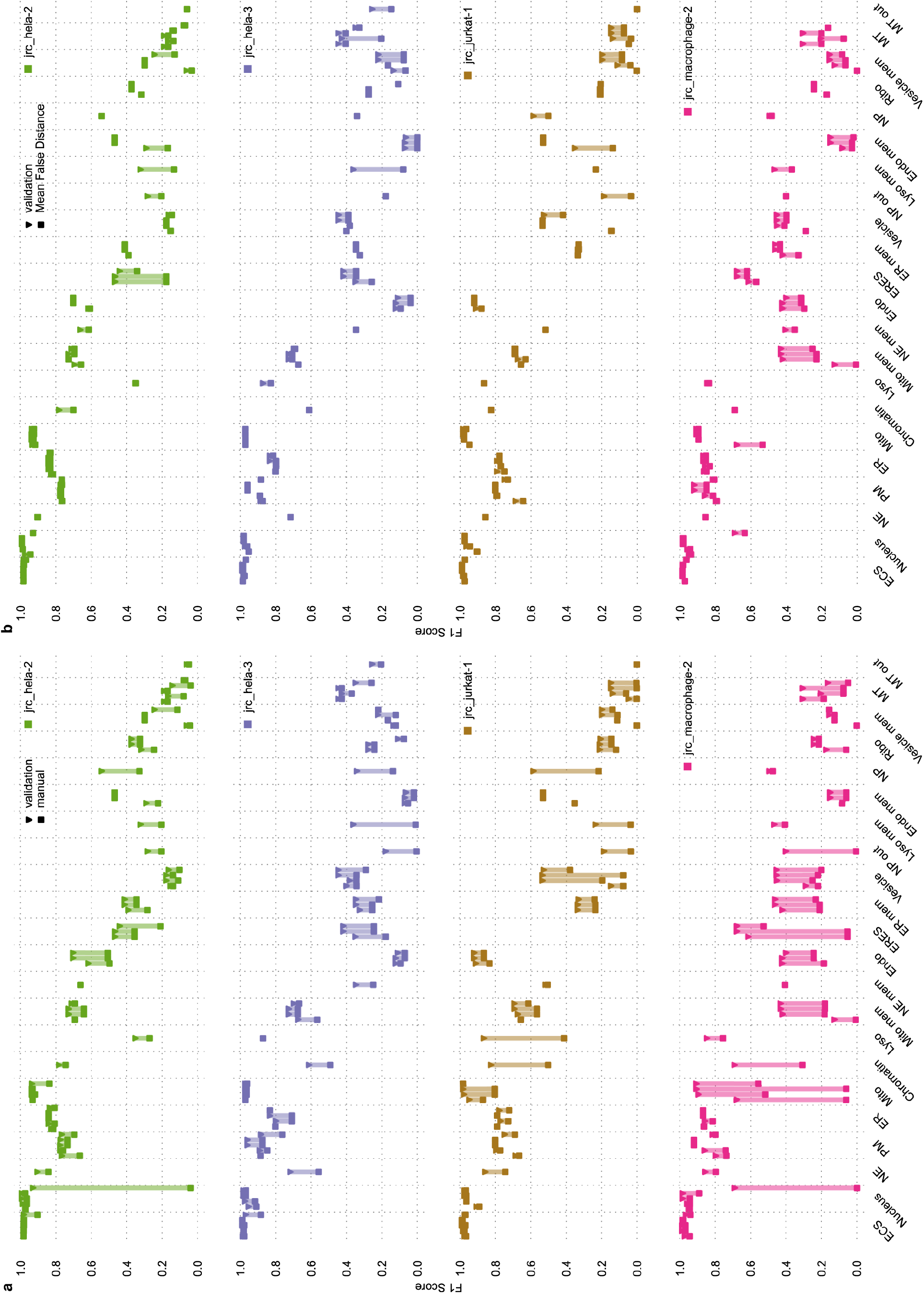
Evaluation metrics & Comparison of manual and automatic hyperparameter tuning. (a) F1 Scores on holdout blocks from four datasets comparing manual and automatic hyperparameter tuning. Data includes all results we collected using the manual comparison of predictions on whole cells, i.e. comparisons across iterations only as well as comparisons across the best iterations of different network types. For automatic validation scores equivalent queries were made against the database. (b) F1 Scores on holdout regions from four datasets comparing F1 Score and Mean False Distance as the metric used for hyperparameter tuning. Data points are equivalent to those from (a).

**Extended Data Fig. 4 -.**
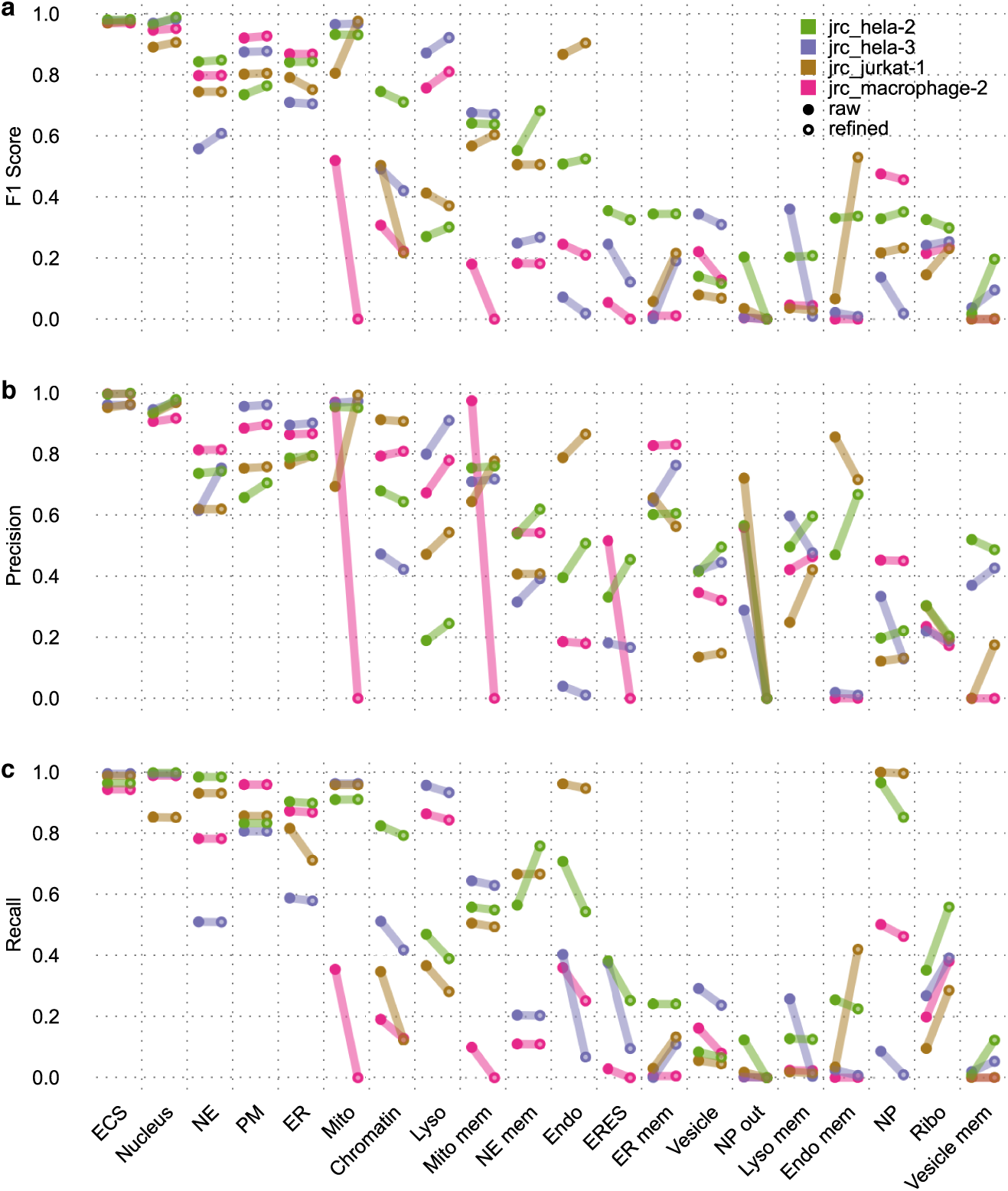
Mitochondria overmerging corrections. (a) Whole cell, 3D rendering of mitochondria in *jrc_hela-2* segmented using naive connected component analysis method. (b) FIB-SEM and raw mitochondria predictions thresholded at 127 (d = 0nm) for the boxed region in (a), shown for one 2D slice. (c) Naive connected component segmentation of mitochondria for the region in (a), performed on smoothed, thresholded predictions at 127 and followed by size filtering and hole-filling. (d) To alleviate over-merging of mitochondria, we smooth the predictions and perform watershed segmentation on all voxels greater than or equal to 127. Shown are the resultant watershed fragments. (e) To create the improved mitochondria segmentations, we agglomerate adjacent fragments in (d) based on parameters that best optimize the resultant segmentations, as chosen by an expert user. (f) Final whole cell rendering of corrected mitochondria predictions.

**Extended Data Fig. 5 -.**
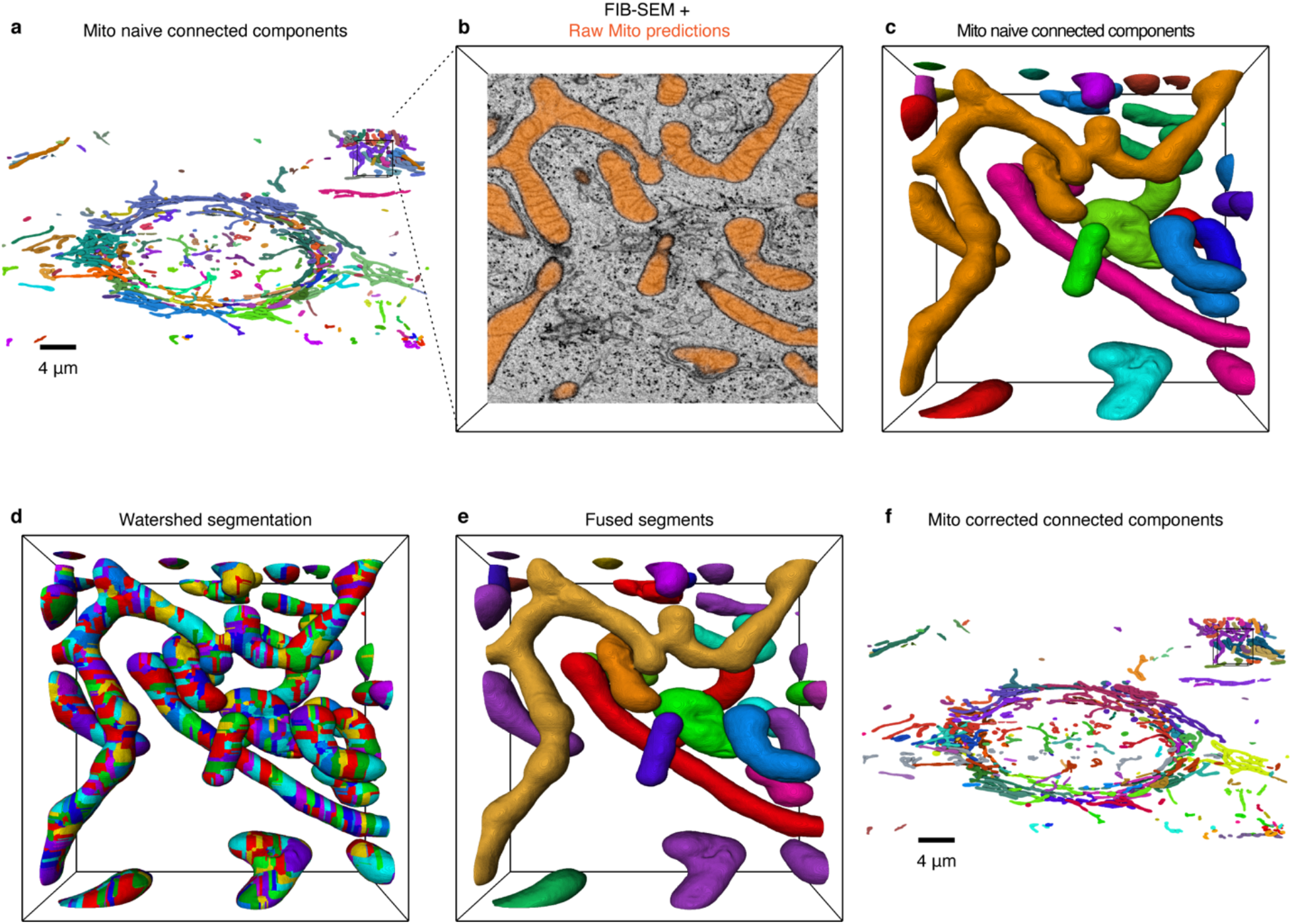
Effect of prediction refinements on evaluation metrics. F1 Score (a), Precision (b) and Recall (c) on holdout blocks from four datasets before (raw) and after (refined) refinements described in Supplementary Methods: Refinements. Network type and iterations represented here are listed in Supplementary Table 1 and are optimized manually with a bias towards potential improvement through the refinement process.

**Extended Data Fig. 6 -.**
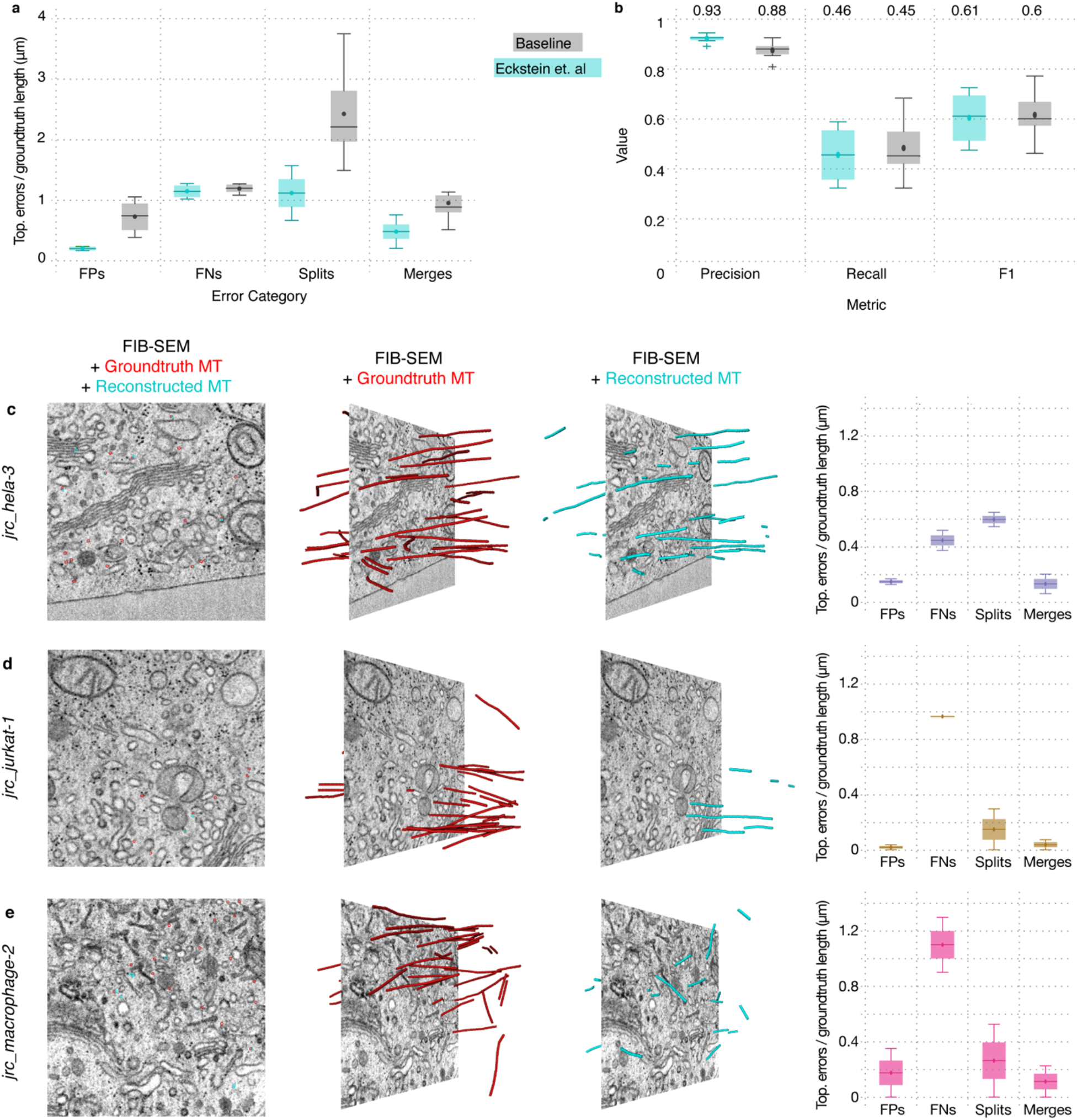
Microtubule Refinement. Comparison of the baseline microtubule refinement and the method described in Eckstein *et al.*^28^. Shown is the accuracy in terms of topological errors on full tracks (a) and precision and recall on individual edges (b), where an edge is correct, if the edge connects two reconstruction vertices that are matched to the same ground truth microtubule track. Each column shows the accuracy of both methods, acquired via 6-fold cross validation over 4 ground truth annotation blocks, where we used two blocks for validation and the remaining two for testing for each run. Numbers above each column in (b) are the median value of the 6 cross-validation runs. See Supplementary Table 4 for a complete listing of evaluation results. 2D FIB-SEM slice with ground truth and reconstructed microtubules in the plane, 3D renderings of the ground truth (red) and reconstructed microtubules (cyan) in selected test blocks on jrc_hela-3 (c), jrc_jurkat-1 (d) and jrc_macrophage-3 (e). Plots show the topological errors normalised by ground-truth microtubule cable length for each cell respectively. For accuracy results in terms of precision, recall and F1 score, see Fig. 2. See Supplementary Table 4 for a complete listing of evaluation results. Standard box plots are used showing the total range (whiskers), outliers (points), the first and third quartile (box), and median (line).

**Extended Data Fig. 7 -.**
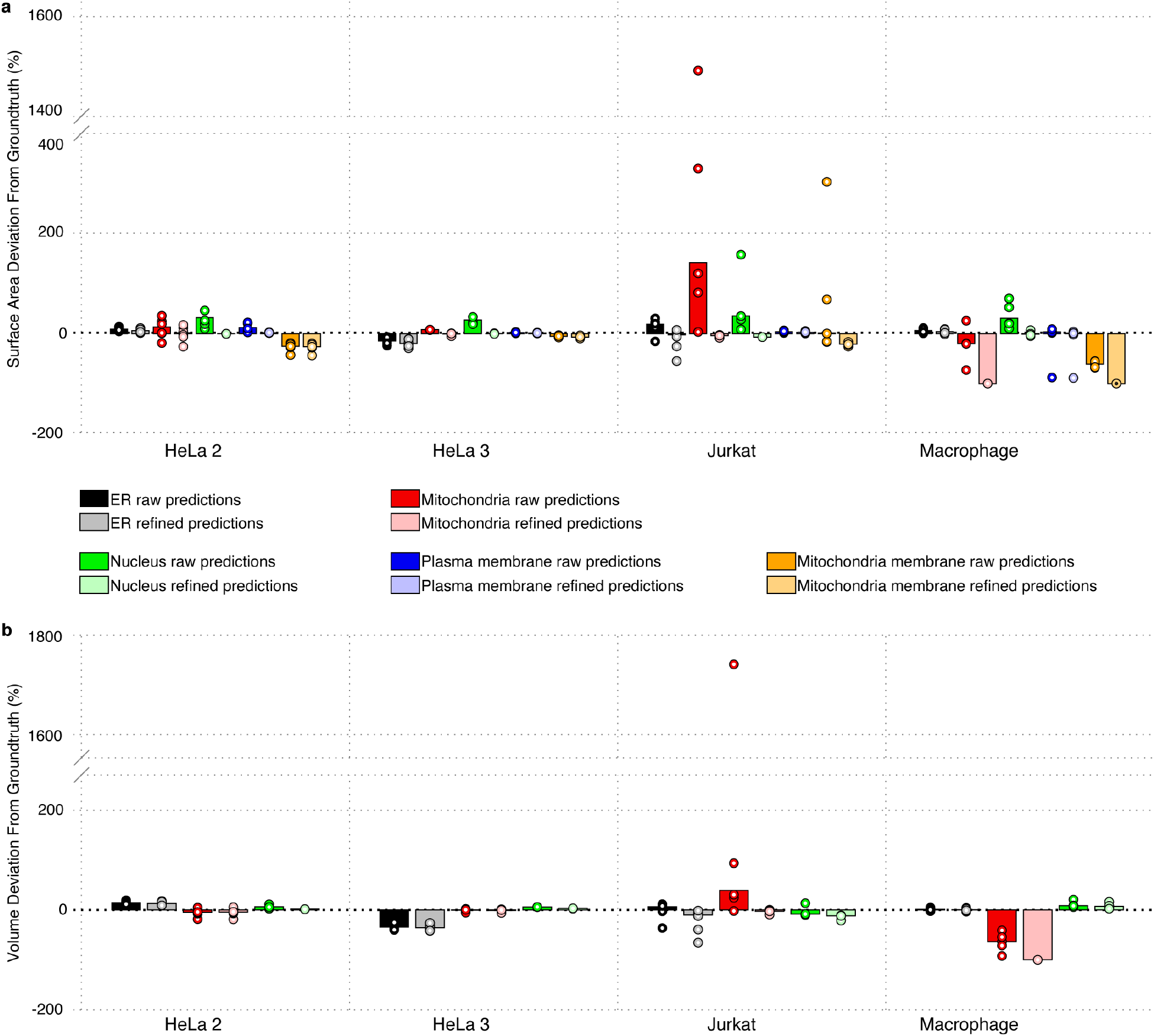
Measurements in holdout regions. Surface area (a) and volume (b) deviations from the ground truth for sample organelles in the holdout regions. Analysis was performed on both raw predictions thresholded at 127 (dark shade), and refined predictions (light shade). In order to get some sense of reliability, we divide each holdout region in half in 3 dimensions resulting in 6 regions. The measurements within these 6 regions are shown (circles), as well as the measurements for the entire holdout region (bar).

**Extended Data Fig. 8 -.**
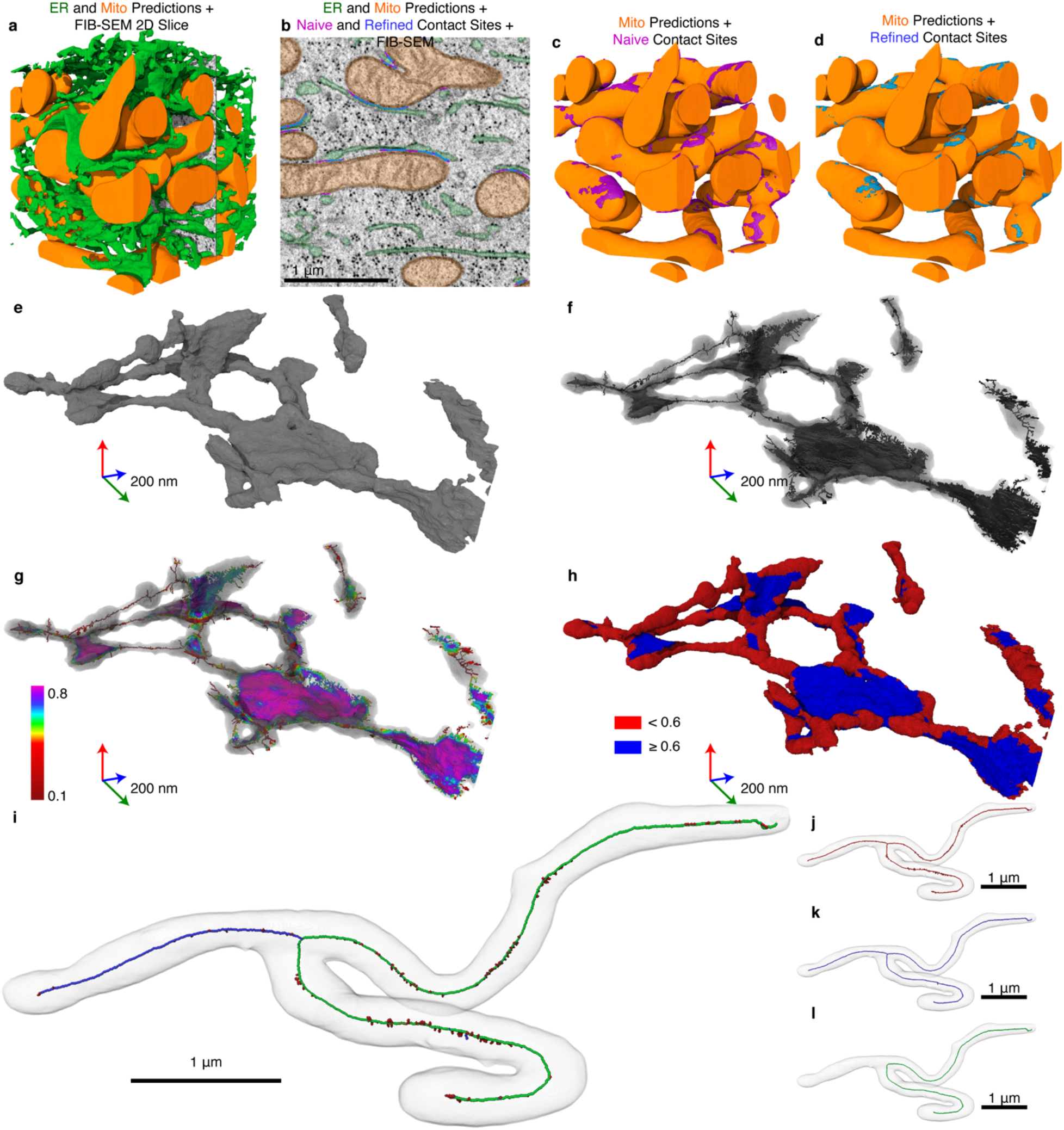
Organelle contact sites, planarity and skeletons. (a) ER (reconstructed, green) and mitochondria (orange) dense regions of *jrc_hela-2* chosen for comparison, with 2D FIB-SEM slice also displayed. (b) 2D FIB-SEM slice shown in (a) displays ER predictions (green), mitochondria predictions (orange), naive contact sites (magenta) and refined contact sites (blue), which are subsets of the naive contact sites. (c) 3D rendering of mitochondria (orange) and simple contact sites (magenta). (d) 3D rendering of mitochondria (orange) and refined contact sites (blue). (e) 3D rendering of refined ER segmentation in an example region of *jrc_hela-2.* (f) Medial surface (black) produced from iterative topological thinning of the ER segmentation (gray). (g) A planar metric (color) is calculated for each voxel in the medial surface based on the ER’s Hessian matrix eigenvalues at that voxel; higher values correspond to more planar regions. (h) The planarity metric medial surface in (c) is used to reconstruct a curvature-labeled ER which is thresholded at 0.6, above which voxels are considered planes (blue) and below which voxels are considered non-planar (red). (i) Topological thinning is used to produce skeletons. Shown is a 3D rendering of an example mitochondria (gray), skeleton (red), pruned skeleton (blue), and longest shortest path (green) from *jrc_hela-2*. (j) Unpruned skeleton used as a starting point. (k) Repetitive pruning produced a final skeleton such that no remaining branch was shorter than 80 nm. (l) Mitochondrial length and average radius were calculated using the longest shortest path within the pruned skeleton. See Supplementary Methods: Quantifications for in depth description.

**Extended Data Fig. 9 -.**
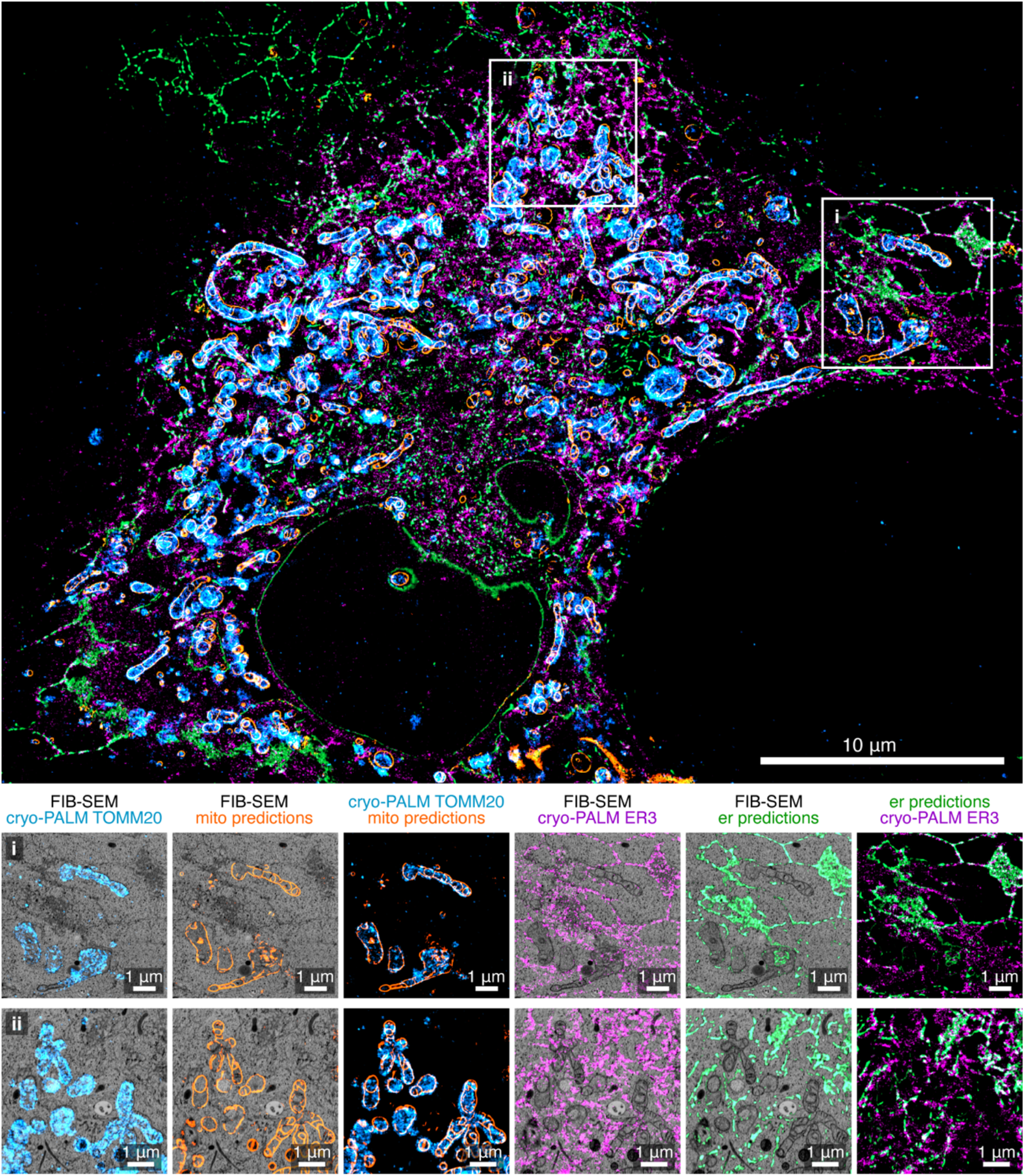
CLEM. Overlays of PALM images and network mitochondria / ER predictions. The substantial similarity between transformed ER predictions and TOMM20 indicates good algorithm performance, despite imperfect staining and false positive / negative predictions. The conclusion is similar when comparing ER predictions and mEmerald-ER3. While the ER predictions have a higher false negative rate than the mitochondria predictions, similar structures overlap after registration.

## Notes

https://openorganelle.janelia.org/

### Competing Interest Statement

The authors have declared no competing interest.

