## Supplementary Information for "Automatic whole cell organelle segmentation in volumetric electron microscopy"

#### Related Works

Innovation in the automatic analysis of volumetric EM has largely been driven by connectomics and thus focused on the segmentation of cells[3–6] and synaptic junctions[7–11] for the reconstruction of the underlying wiring diagram. In addition, mitochondria, which present a very salient structure in these data, are often treated as an alternative segmentation task[9,12–19]. Studies on other organelles[20,21] and other tissue or cell types[22,23] have been less frequent. Recent work has included the segmentation of the nuclear envelope in HeLa cells[24,25], detection of mitochondria and endolysosomes in urothelial FIB-SEM data[26], the joint reconstruction of mitochondria and endoplasmic reticulum in serial section EM data of mouse cortex[27] andthe tracking of microtubules in ssTEM data[28].

A major hurdle for the advancement of automated methods for other organelles and data types is the need for large amounts of training data. This is particularly the case for deep learning based methods[9,16,18,26] that - following a general pattern in computer vision[57] - have now replaced more traditional computer vision approaches[12,14,15,17] as the state of the art.

The vast size of connectomic datasets often warrants expensive annotation of dedicated training data. Several challenges have provided publicly available training data[58–60] which has enabled a broader research community to work on cell and synapse segmentation. Several ground truth datasets for mitochondria segmentation are also freely available[12,13,18–20,26]. Any one ground truth dataset however is unlikely to facilitate inference in different domains. A few studies have started to explicitly address this problem through domain adaptation, i.e. the ability of a classifier that has been trained on one domain to be transferred to a different domain where training data is scarce or non-existent. Haberl et al.[61] finetune a network trained on serial section data to predict membranes in block face SEM. Bermundez-Chacon *et al.*[62] train a two-stream U-Net architecture where some of the weights are shared between the two domains which represent two distinct areas of a mouse brain. Roels *et al*.[63] test strategies for aligning the feature space learned for the source and target domain, either through an additional feature similarity loss term or an auxiliary training task of reconstructing the inputs. They validate on mitochondria segmentation in tissue imaged with ssTEM transferred to HeLa cells imaged with FIB-SEM.

In practice, many biologists rely on commercial, closed source solutions[64] or semi-automatic workflows for segmentation, as facilitated by tools such as Amira-Avizo, Imaris[65], dragonfly[66], ilastik[67] or MIB[68]. However, these tools involve substantial user interaction and often significant expertise to properly segment challenging datasets. Notably, Müller et al.[69] have developed a set of specialized automatic, semi-automatic, and manual methods to reconstruct mitochondria, the Golgi apparatus, centrioles, insulin secretory granules, and microtubules in seven beta cells contained in two large FIB-SEM volumes.

### Supplementary Methods

#### Datasets

Below is a brief description for each sample including cell type, general sample preparation details, and original publication.

***jrc_hela-1***

*Description*: Wild-type, interphase HeLa cell (ATCC CCL-2).
*Protocol*: High pressure freezing, freeze-substitution resin embedding with 2% $OsO_4$ 0.1% UA 3% $H_2O$ in acetone; resin embedding in Eponate 12.
*Contributions*: Sample provided by Aubrey Weigel (HHMI/Janelia), prepared for imaging by Gleb Shtengel (HHMI/Janelia), with imaging and post-processing by C. Shan Xu (HHMI/Janelia).
*Publication*: unpublished
*Voxel size*: 8 nm × 8 nm × 8 nm
*OpenOrganelle link*: https://openorganelle.janelia.org/datasets/jrc_hela-1
*EM data DOI:* 10.25378/janelia.13123415
*Segmentation DOI:* 10.25378/janelia.13120280

***jrc_hela-2***

*Description*: Wild-type, interphase HeLa cell (ATCC CCL-2).
*Protocol*: High pressure freezing, freeze-substitution resin embedding with 2% $OsO_4$ 0.1% UA 3% $H_2O$ in acetone; resin embedding in Eponate 12.
*Contributions*: Sample provided by Aubrey Weigel (HHMI/Janelia), prepared for imaging by Gleb Shtengel (HHMI/Janelia), with imaging and post-processing by C. Shan Xu (HHMI/Janelia).
*Publication*: Xu et al., 2020[2]
*Voxel size*: 4 nm × 4 nm × 4 nm
*OpenOrganelle link*: https://openorganelle.janelia.org/datasets/jrc_hela-2
*EM data DOI:* 10.25378/janelia.13114211
*Segmentation DOI:* 10.25378/janelia.13108343

***jrc_hela-3***

*Description*: Wild-type, interphase HeLa cell (ATCC CCL-2).
*Protocol*: High pressure freezing, freeze-substitution resin embedding with 2% $OsO_4$ 0.1% UA 3% $H_2O$ in acetone; resin embedding in Eponate 12.
*Contributions*: Sample provided by Aubrey Weigel (HHMI/Janelia), prepared for imaging by Gleb Shtengel (HHMI/Janelia), with imaging and post-processing by C. Shan Xu (HHMI/Janelia).
*Publication*: Xu et al., 2020[2]
*Voxel size*: 4 nm × 4 nm × 4 nm
*OpenOrganelle link*: https://openorganelle.janelia.org/datasets/jrc_hela-3
*EM data DOI:* 10.25378/janelia.13114244

*Segmentation DOI:* 10.25378/janelia.13117586

***jrc_jurkat-1***

*Description*: Wild-type Jurkats. Clone E6-1 (ATCC TIB-152).
*Protocol*: High pressure freezing, freeze-substitution resin embedding with 2% $OsO_4$ 0.1% UA 3% $H_2O$ in acetone; resin embedding in Eponate 12.
*Contributions*: Sample provided by Huxley Hoffman and Schuyler van Engelenburg (U. Denver), prepared for imaging by Gleb Shtengel (HHMI/Janelia), with imaging and post-processing by C. Shan Xu (HHMI/Janelia).
*Publication*: Xu et al., 2020[2]
*Voxel size*: 4 nm × 4 nm × 4 nm
*OpenOrganelle link*: https://openorganelle.janelia.org/datasets/jrc_jurkat-1
*EM data DOI:* 10.25378/janelia.13114259
*Segmentation DOI:* 10.25378/janelia.13117697

***jrc_macrophage-2***

*Description*: Wild-type THP-1 macrophage. THP-1 human monocyte cell line (ATCC TIB-202) treated with PMA to differentiate into macrophages.
*Protocol*: High pressure freezing, freeze-substitution resin embedding with 2% $OsO_4$ 0.1% UA 3% $H_2O$ in acetone; resin embedding in Eponate 12.
*Contributions*: Sample provided by Huxley Hoffman and Schuyler van Engelenburg (U. Denver), prepared for imaging by Gleb Shtengel (HHMI/Janelia), with imaging and post-processing by C. Shan Xu (HHMI/Janelia).
*Publication*: Xu et al., 2020[2]
*Voxel size*: 4 nm × 4 nm × 4 nm
*OpenOrganelle link*: https://openorganelle.janelia.org/datasets/jrc_macrophage-2
*EM data DOI:* 10.25378/janelia.13114343
*Segmentation DOI:* 10.25378/janelia.13117745

***jrc_sum159-1***

*Description*: Wild-type SUM-159 cell, treated with 0.5 mM oleic acid for 45 mins prior to high pressure freezing to induce the formation of lipid droplets.
*Protocol*: High pressure freezing, freeze-substitution resin embedding with 2% $OsO_4$ 0.1% UA 3% $H_2O$ in acetone; resin embedding in Eponate 12.
*Contributions*: Sample provided by Jeeyun Chung, Tobias Walther and Bob Farese (Harvard U.), prepared for imaging by Gleb Shtengel (HHMI/Janelia), with imaging and post-processing by C. Shan Xu (HHMI/Janelia).
*Publication*: Xu et al., 2020[2]
*Voxel size*: 4 nm × 4 nm × 4 nm
*OpenOrganelle link*: https://openorganelle.janelia.org/datasets/jrc_sum159-1
*EM data DOI:* 10.25378/janelia.13114352
*Segmentation DOI:* 10.25378/janelia.13118156 (manual annotations only)

***jrc_choroid-plexus-2***

*Description*: Mouse choroid plexus
*Protocol*: Postfixed in 1.0% osmium tetroxide in 0.1M cacodylate buffer (pH 7.4) for 1 hour at room temperature. Following postfixation, the samples were rinsed with buffer, dehydrated through a graded series of ethanol, and embedded in durcupan.
*Contributions*: Sample provided by Christopher Walsh (Harvard), prepared for imaging by Song Pang (HHMI/Janelia), with imaging by Song Pang (HHMI/Janelia) and C. Shan Xu (HHMI/Janelia), and post-processing by C. Shan Xu (HHMI/Janelia).
*Publication*: Coulter et al., 2018[38], Xu et al., 2017[40]
*Voxel size*: 8 nm × 8 nm × 8 nm
*OpenOrganelle link*: https://openorganelle.janelia.org/datasets/jrc_choroid-plexus-2
*EM data DOI:* 10.25378/janelia.13123427
*Segmentation DOI:* 10.25378/janelia.13122509

***jrc_cos7-11***

*Description*: COS-7 cell overexpressing mEmerald-ER3 and Halo/JF525-TOMM20 (ATCC CRL-1651)
*Protocol*: High pressure freezing, freeze-substitution resin embedding with 2% $OsO_4$ 0.1% UA 3% $H_2O$ in acetone; resin embedding in Eponate 12.
*Contributions*: Sample provided by Melanie Freeman (UC Berkley), prepared for imaging by Gleb Shtengel (HHMI/Janelia), with imaging and post-processing by C. Shan Xu (HHMI/Janelia).
*Publication*: Hoffman et al., 2020[39]
*Voxel size*: 8 nm × 8 nm × 8 nm
*OpenOrganelle link*: https://openorganelle.janelia.org/datasets/jrc_cos7-11
*EM data DOI:* 10.25378/janelia.13123385
*Segmentation DOI:* 10.25378/janelia.13120265

#### Data Pre-Processing

To create image volumes used for machine learning, raw planes from the FIB-SEM microscope were aligned to create a volume, then resampled at nearly isotropic resolution; Voxel values were then converted to unsigned 8 bit data type. See Xu *et al*.[2,40] for more details. We manually adapted the contrast of each dataset to be as consistent as possible with other datasets. Volumes were stored in a chunked array file format in N5 containers, which facilitated performant random access to different regions of the imaging volume[42].

FIB-SEM samples are embedded in resin, which can be distinguished from cells and tissue by relatively simple image filter routines. Because we know in advance that there are no subcellular structures of interest outside the cell, we would like to exclude resinous regions from computationally expensive steps like model inference. Thus, for each volume used for machine learning, we created a binary mask to separate resin from sample material. Masks were created by applying an image filter that responds to local disorder, such as local entropy or standard deviation filters, then applying Gaussian blur and thresholding. For each volume, the threshold was tuned to include all of the sample of interest, at the expense of including a small amount of marginal resin.

#### Training Data

Manual segmentation was performed on acquired 4 nm × 4 nm × 4 nm resolution datasets of HeLa, Jurkat, Macrophage, and SUM159 cells using 3D visualization and processing software platforms Amira-Avizo (ThermoFisher) and BigCat[44]. Cropped and stacked TIF files, often 0.5 $\mu m^3$ in volume, were imported to Amira to be annotated on orthogonal planes XY, YZ, and XZ. The source TIF file voxel size was artificially upsampled to 2 nm × 2 nm × 2 nm, output type was assigned as 8-bit label, and a variety of image processing filters were applied to increase image clarity, contrast, and smoothness based on FIB-SEM staining variation. Expert annotators used numerous selection tools e.g. brush, interpolation, fill, lasso, and smooth to annotate and assign labels to all voxels. The resulting 3D rendering was used as a contextual guide for further segmentation and organelle identification.

For additional spatial and morphological context, especially concerning structures along the edge of each stack, label fields were exported as 3D TIF files and converted to HDF5 format for visualization and annotation in BigCat. A padding of ~200 voxels was added to each block to visualize surrounding cellular context and reduce ambiguity. Rough BigCat annotations were saved, converted back to TIF format, and imported into Amira for further annotation. Upon completion, each label field was consolidated into a final label field, each Amira 'material' representing an organelle class. Finally, cross-annotator quality checks ensured every voxel was annotated, material texture was smoothed, cleaned, properly identified and separated from neighboring materials. In preparation for model training, blocks were stored as chunked arrays in an N5 container and metadata for each block were stored in a database.

#### Organelle Classification

Organelles were manually identified using morphological features established in the literature[43]. Because every voxel within an annotated block must be classified, 'hollow' organelles e.g. endoplasmic reticulum, which contains lumen bound by a membrane, are defined by 'lumen' (lum) and 'membrane' (mem), or 'inside' (in) and 'outside' (out) depending on organelle composition. Below is a glossary of the classes including their proper name, the shorthand notation used throughout the manuscript, related subclasses when applicable, and a brief description. Examples of each class can be found in Extended Data Fig. 1.

*Long name (shorthand notation of classes included if superclass)*: Description.

*Actin*: Cytoskeletal filaments that are often found in globular patches throughout the cytosol. Actin is usually lightly stained and appears in dynamic, wavy clusters. Unlike microtubules, actin filaments do not have a visible diameter and have unrestricted curvature. Actin is not currently an object class and was only included as negative training examples.

*Centrosome (Centrosome, Centrosome D App, Centrosome SD App)*: Barrel-shaped structure composed of microtubule triplets. Centrioles are often found in pairs and microtubule staining is dark and distinct. A cross section of centriole ends is a round, nine-fold star shape. Skeleton annotations in BigCat were used to trace each microtubule of the barrel structure. These skeletons were then used to inpaint a full microtubule triplet into the volume. Voxel classification was used to annotate distal (D App) and subdistal appendages (SD App).

*Cytosol*: Defined as any volume enclosed by a plasma membrane that is not categorized within an organelle or molecule class. Cytosol is often identified as the negative space within a cell surrounding relatively strongly stained organelles, proteins, and other molecules. The cytosol was not used as a training class, but rather as a negative example for all other training classes.

*Endosomal Network (Endo, Endo mem, Endo lum)*: Light lumen organelles that constitute the endosomal network. These structures have a few characteristic morphologies, including ‘ribbon’ structures and spheres with multiple membrane invaginations. The endosomal network class includes autophagosomes, endosomes, multivesicular bodies, and peroxisomes as EM staining alone is not sufficient for differentiation.

*Endoplasmic reticulum (ER, ER mem, ER lum)*: An extensive network of tubular structures often studded with ribosomes. The endoplasmic reticulum is distinct from multivesicular bodies based on connectivity; ER networks always connect back to themselves and ultimately connect back to the nuclear envelope. In contrast, the morphologically-similar multivesicular bodies are disconnected. ER exit sites (ERES), nuclear envelope (NE), and associated nuclear pores (NP) are included in the ER superclass.

*Endoplasmic reticulum exit sites (ERES, ERES mem, ERES lum)*: A cluster of tubular structures and vesicles that bud from the endoplasmic reticulum. Endoplasmic reticulum exit site lumen and morphology is generally consistent with the corresponding ER network.

*Euchromatin (E Chrom, N-E Chrom)*: Light, single chromatin within the nucleus. Euchromatin stain is lighter and less compact than heterochromatin. Nucleolus euchromatin (N-E Chrom) is not associated with or connected to euchromatin (E Chrom) outside the nucleolus.

*Extracellular Space (ECS)*: Defined as any volume outside of the plasma membrane boundary, including both cellular plasma membrane and membrane surrounding secreted vesicles; it does not contain any stained organelles or molecules and is not defined by a particular morphology or contrast.

*Golgi (Golgi, Golgi mem, Golgi lum)*: Stacked, ‘pancake-like’ structures that uniformly morph and bend together. Gaps of a similar thickness to each Golgi layer compose a Golgi apparatus that often consists of 5-7 layers. Stacks may be interconnected and often connect to ER network and endosomal network (Endo). Surrounding and budding vesicles are often included in the Golgi network.

*Heterochromatin (H Chrom, N-H Chrom)*: Dark clusters of chromatin within the nucleus. Heterochromatin stain is darker and more compact than euchromatin. Nucleolus heterochromatin (N-H Chrom) is not associated with or connected to heterochromatin (H Chrom) outside the nucleolus.

*Lipid Droplets (LD, LD mem, LD lumen)*: Spherical organelles enclosed by a lipid monolayer and characterized by a shriveled, 'lumpy' morphology due to general staining. Lipid droplets (LD) are generally lighter than surrounding cytosol and have subtle membrane staining.

*Lysosome (Lyso, Lyso mem, Lyso lum)*: Spherical, dark lumen organelles that constitute the late endosomal network. Lysosomes can have multiple membranes; the lysosome class also includes autophagosomes, multivesicular bodies, endosomes, and peroxisomes as EM staining alone is not sufficient for differentiation.

*Microtubules (MT, MT out, MT in)*: Cylindrical, cytoskeletal polymers characterized by restricted curvature and a 25 nm diameter. Microtubules often run parallel, do not branch, and may appear to pierce through other organelles, especially ER.

*Mitochondria (Mito, Mito mem, Mito lum, Mito Ribo*): Large, ovoid organelles characterized by outer and inner membranes that form cristae. Mitochondria can fuse and branch to form tubular networks. Inner membrane folding density varies based on cell type. Dark-staining aggregates within mitochondria lumen have been identified and termed mitochondrial ribosomes (Mito Ribo).

*Nuclear envelope (NE, NE mem, NE lum)*: An extensive membrane-like structure identified by the presence of two lipid bilayers often studded with ribosomes. The nuclear envelope connects back to itself and is continuous with ER networks. The double membrane structure is perforated with nuclear pores (NP) and establishes a boundary between chromatin (Chrom) and the cytosol.

*Nuclear pores (NP, NP out, NP in)*: Circular, 120 nm pores in the nuclear envelope. When viewing a cross section of the nucleus, nuclear pores appear as breaks or gaps in envelope connectivity. Nuclear pores span both bilayers of the nuclear envelope (NE).

Nucleolus *(Nucleolus, N-E Chrom, N-H Chrom)*: A dark, dense spherical structure within the nucleus containing heterochromatin (N-H Chrom) and euchromatin (N-E Chrom).

*Nucleus (Nucleus, E Chrom, N-E Chrom, H Chrom, N-H Chrom, NE memb, NE lum, NP out, NP in, Nucleolus, Nucleoplasm)*: The largest spherical organelle characterized by patches of high contrast chromatin and a dark, central nucleolus. In addition to heterochromatin and euchromatin, the nucleus class includes the nuclear envelope and associated nuclear pores, the nucleolus and associated nucleolus heterochromatin and euchromatin, and surrounding nucleoplasm. Our framework allows for a mix of annotating the nucleus via its subclasses or with a generic nucleus label.

*Plasma membrane (PM)*: A thin, extensive bilayer that surrounds the cell and separates extracellular space from cytosol. To be classified as plasma membrane, the membrane must always connect back to itself. Otherwise, the membrane is most likely part of the endosomal network (Endo).

*Ribosomes (Ribo)*: Macromolecular structures characterized by darkly stained ellipsoid spots which result from diffraction-limited resolution. Ribosomes are often found in clusters bound to ER or suspended in cytosol. Point annotations in BigCat were used to demark the centroid of ribosomes in a volume. These were then painted into a separate volume as points and virtually expanded to 18 nm as needed.

*Vesicle (Vesicle, Vesicle mem, Vesicle lum)*: Small, spherical organelles less than 100 nm in diameter with lumen varying in color depending on protein content. Vesicles are often found in clusters surrounding Golgi and ER.

#### Machine Learning

We used 3D U-Net architectures[29,30], a type of architecture that has been very successful for this type of computer vision problem. On each level we used 2 convolutional layers with kernel size (3, 3, 3), valid padding and rectified linear units. Our architectures have 4 levels with 3 downsampling layers with factors (2, 2, 2), (3, 3, 3) and (3, 3, 3) between them. Starting from an initial feature width of 12, we multiplied that number by a factor of 6 with each downsampling operation. For upsampling we used a transpose convolutional layer with a kernel that is a constant. In order to enforce translational equivariance, we used valid padding and feature maps were cropped to a multiple of the cumulative upsampling factor after each upsampling operation. Due to the large number of output maps in some networks we didn't decrease the number of features back to 12 for the final two convolutions. To obtain the correct number of outputs we used a convolutional layer with kernel size (1, 1, 1). Networks were trained with Tensorflow[70] in Gunpowder[71].

In addition to this more standard 3D U-Net architecture that we fed with raw data of approximately 4 nm isotropic resolution and that outputs images at the same resolution, we also trained networks that increase the resolution from 8 nm isotropic input to 4 nm isotropic output. These U-Nets had the same hyperparameters as described above but with an additional level without a skip connection after upsampling with a factor of (2, 2, 2). Again, we kept the number of feature maps at 72 until reducing it to the desired number of output layers. To simulate 8 nm isotropic raw data from our existing set of training data that had been imaged at approximately 4 nm isotropic resolution, we subsampled the image data by picking a random voxel from each block of (2 × 2 × 2) voxels. This downsampling scheme should have a more realistic noise structure than downsampling by averaging voxels.

The loss was computed as the sum of the mean squared error between each output map and the signed tanh distance transform of each label[10]. All experiments scaled the distance transform by a factor 50.

The loss was minimized using the Adam optimizer[45] with a learning rate of $5 \times 10^{-5}$. The loss for each class was balanced to give equal weight to the positive and negative labels in a patch. The fraction of positive voxels used to calculate the weighting factors was cut off to be between 0.05 and 0.95. For the subclasses we typically used the weighting matrix of the superclass.

The distance transforms were computed on-the-fly from the ground truth annotated at (2 nm × 2 nm × 2 nm) resolution. They were computed on patches sufficiently larger than the output size of the network and are then cropped to the network output size and downsampled to (4 nm × 4 nm × 4 nm). This guarantees the correctness of the distance transforms on the full output volume after the application of our augmentation procedures but is computationally very expensive. Parts of the patch where correctness cannot be guaranteed due to block boundary effects are not considered for loss computation. For ribosomes and centrosomes, which were annotated with point and line annotations converted into a tube structure, respectively, we added a constant to the distance transforms computed from those annotations to reflect their true size.

Setups jointly trained for different numbers of classes were used to examine the optimal multi-class strategy. Type “all” networks train all classes jointly, type “common” networks train the 14 classes with instances in at least 10 training blocks. Lastly, type “few” networks specialize in one or two organelles, encompassing up to 4 classes. The type “all” network was only trained for (4 nm × 4 nm × 4 nm) input data. In some networks we tried to slightly reduce the number of classes by not including redundant ones, i.e. classes that can be computed using the other classes trained by the network. Supplementary Table 1 summarizes all network types.

During training patches were sampled randomly from all training blocks with the probability for each proportional to its size. Patches needed to have at least 35,000 voxels annotated to be processed for training. Some type “few” networks that are specialized on labels that have very few positive labels in our training set failed to learn anything using this sampling procedure. For ribosomes, vesicles and golgi we instead sampled with probability 0.5 from the set of training blocks containing positive annotations and then using our standard sampling procedure within that set. This adaptation was only made for the 4 nm networks.

To virtually increase the size of our training set and improve robustness to contrast variations, each patch was augmented. The augmentations included random flips and rotations, elastic deformations and linear as well as gamma intensity augmentations. For the 8 nm networks we chose one of 8 pre-generated versions of the randomly downsampled raw data at random which can be interpreted as an augmentation mimicking different instances of shot noise.

Primarily due to the on-the-fly computation of the distance transform, the training time for the presented networks is substantial. We roughly estimated that on average one training iteration took between 2.1 s to 5.2 s for type “few” networks, 5.4 s for type “many” networks and 12 s for type “all” networks with large variability in all cases. The 8 nm networks take ~3 - 3.5 times longer to train than the corresponding 4 nm networks due to the larger output sizes. This training time could most likely be substantially reduced through both optimizing the implementation as well as allowing a larger number of CPUs to supply training examples to one GPU, which is currently fixed in our setup where type “many” and “all” networks used 12 CPUs and 1 Tesla 32 GB V100 card and type “few” networks used 5 CPUs and 1 GeForceRTX2080Ti card.

#### Inferences

Inference on whole cells can be performed in parallel by dividing the output volume into non-overlapping chunks as all networks use valid padding and are translationally equivariant. The corresponding chunks of raw data were extended by the field-of-view of the network. The size of input and output blocks can be increased for inference as the activations do not need to be stored for backpropagation. Finally, each output block was converted to uint8 and written to disk in the N5 format[42]. We used Dask[72] to load, preprocess and write chunks while the GPU performs the computationally expensive neural network inference. Depending on the network architecture we achieved inference speeds of 0.5 - 4.8 $\frac{\mathrm{M}vx \cdot \#channels}{s \cdot GPU}$ on GeForceRTX2080Ti cards, which provide 11 GB of GPU RAM and 12 TFLOPS. To save computational costs as well as storage space we restricted inference to the foreground masks described above. We evaluated networks at intervals of 25k iterations for manual and automatic validation.

#### Evaluations

*Manual Validation*

A custom FIJI plugin was created for manual evaluation of predictions to determine which was best for each organelle within a dataset. In the plugin, the user selects two predictions to compare as well as an organelle class to evaluate. A random 150 × 150 × 150 voxel crop is then chosen from the volume and cropped from each prediction. If neither prediction contains at least 500 organelle voxels (those with signed tanh distance transform ≥ 0, corresponding to voxel values ≥ 127), a new random crop is selected. The selected cropped predictions are shown to the user as two image stacks, with the predictions thresholded at 127 and overlaid on the raw data. The user inspects the stacks and selects which prediction - if any - did a better job of predicting the chosen organelle. The crops are randomly labeled so that the user is blind to which image came from which prediction. Once a crop has been evaluated, the plugin calculates and reports the p-value corresponding to the null hypothesis that the two networks perform equally well. This is analogous to determining if a coin is unbiased and corresponds to a binomial distribution B(n, p) with n trials and probability of success p = 0.5. This evaluation process then repeats for a new crop.

100 such evaluations were performed. If the p-value < 0.01 after the 100 evaluations, we rejected the null hypothesis and the network that performed best was deemed the winner. Otherwise, 50 more crops were evaluated. At that point if p-value > 0.01, a winning network was chosen based on visual inspection using N5-viewer. No ties were allowed.

To determine the optimal network for each organelle class using this method, the user first compared predictions from the same network but different iterations. For each network, a range of 200k iterations were evaluated in this manner. To establish this range, the user performed a visual inspection using N5-viewer in order to determine the approximate 200k range of iterations to be evaluated via the UI. Once the best iteration for each network had been selected, the user compared the best iteration predictions from different networks using this method to find the optimal network and iteration combination.

*Validation with ground truth*

To validate the manual evaluation method we used holdout blocks from four datasets (*jrc_hela-2, jrc_hela-3, jrc_macrophage-2* and *jrc_jurkat-1*) to quantitatively compare the thresholded predictions against the ground truth. We evaluated the following metrics for each label contained in those blocks at intervals of 25k iterations and organized all results in a database. We did not observe that any of our more general conclusions for comparisons between different types of network depended on our choice of metric.

*Precision:*

Precision measures the proportion of (positive) detections that are in fact correct. For samples without detections we define it to be 0.

$$precision = \frac{true\ positives}{true\ positives + false\ positives}$$

*Recall:*

Recall measures the ability of a classifier to find all instances of a class.

$$recall = \frac{true\ positives}{true\ positives + false\ negatives}$$

*F1 Score:*

The harmonic mean of recall and precision is known as F1 score, weighing the two metrics equally. In the context of segmentation it is also often referred to as Dice coefficient. For samples without detections we define it to be 0. The F1 Score is very sensitive to the exact localization of a detection which is often punitive for small or thin objects.

$$F1 = 2 \cdot \frac{precision \cdot recall}{precision + recall}$$

*Mean False Distance (MFD)[60]:*

The Mean False Distance is the arithmetic mean of the average distance of a detection to a positive ground truth annotation and the average distance of a ground truth annotation to a detection. For samples without detections it is not defined. It is designed to be more forgiving with regards to the exact localization than the more standard F1 score and more representative of the effect errors have on many subsequent analyses. It is closely related to the target function we used for training our neural networks.

$$MFD = \frac{1}{2}(MFPD + MFND)$$

with Mean False Positive Distance (MFPD) and Mean False Negative Distance (MFND) defined as

$$MFPD = \mathrm{mean}\Big(\mathrm{dt}(ground\ \ truth)\big|_{prediction>0}\Big)$$

$$MFND = \mathrm{mean}\Big(\mathrm{dt}(prediction)\big|_{ground\ \ truth>0}\Big)$$

, where $\mathrm{dt}$ is the Euclidean distance transform.

Those results allowed us to report validation scores, i.e. optimizing the metric on the block it is being evaluated on, and test scores, i.e. optimizing the mean of the results across all other blocks for each label and dataset. With few exceptions, networks have been trained at least until they started overfitting which we defined as validation scores not improving for at least two checkpoints (50k iterations). We used the same criterion for the lower bound on iterations to evaluate.

Extended Data Fig. 3a shows how automatic evaluation compares to manual evaluation by matching the validation F1 score against the F1 score of the network/iteration chosen through manual validation. These data include both the optimization of only the iteration as well as the comparison across setups as described above. In some cases we observed large discrepancies between the automatic and manual validation score, e.g. for the mitochondria segmentation in *jrc_macrophage-2*. We suppose that those discrepancies result from the block not being representative of the dataset as a whole for that class due to its limited size. Generally, automatic and manual validation agree relatively well. As a guideline for what level of agreement to expect in Extended Data Fig. 3b we compare the validation F1 score as before against the F1 score of the network/iteration chosen by optimizing a different metric, in this case the Mean False Distance.

The manual validation is not limited to the classes appearing in the holdout block and is more representative of the quality of segmentation throughout the dataset. The overall good agreement between automatic and manual validation indicates that the biases introduced by the evaluator having to weigh different kinds of errors is not detrimental. For these reasons we present segmentations obtained with settings optimized manually unless otherwise noted. The corresponding scores are summarized in Supplementary Table 2.

#### Refinements

In general, we wanted to segment the datasets such that every voxel is assigned a label based on whether it is background or part of an organelle. Background voxels were assigned a value of 0, and organelle voxels were assigned IDs that are unique to each individual organelle instance.

This was accomplished by processing and refining the predictions. A description of each type of processing refinement can be found below, and Extended Data Table 1 denotes which refinements were used for each organelle class.

*Smoothing*: A Gaussian filter (σ = 12 nm) was applied to the initial predictions to smooth them out and reduce noise.

*Connected components:* Except for ribosomes and microtubules (discussed later), all organelle predictions were thresholded at a predicted distance of 0 nm, above which voxels were considered part of an organelle; this corresponds to signed tanh distance transform ≥ 0. Connected component analysis of these organelle voxels was then performed to group the voxels into individual organelles with distinct IDs.

*Size Filtering*: Often, a minimum size filter was applied to the connected components in order to eliminate small false positives. The size of the filter was conservative and based on the expected organelle size.

*Hole Filling*: A hole is defined as a region of background voxels completely surrounded by a single organelle. Hole filling was performed by relabeling the hole voxels with the ID of the surrounding organelle.

*Custom Filtering*: When size filtering alone was insufficient, an expert user inspects the results and selects which individual objects should be removed.

*Watershed/Agglomeration:* To prevent overmerging, a combination of watershed segmentation and agglomeration was applied[31,32] (Extended Data Fig.4). This broke up objects that would otherwise be merged under the default connected components analysis. Predictions were first smoothed with a Gaussian (σ = 12 nm), followed by setting all background voxels (signed tanh distance transform < 0 ) to 0. Watershed segmentation was then performed on the resultant volume to oversegment the prediction. This was followed by agglomeration in which adjacent segments were merged if a given percentage of voxels along their shared edge had a predicted distance above a given threshold. The best percentage and threshold were manually chosen based on an expert user evaluating results from a range of percentages and thresholds and choosing those which produced optimal segmentations.

*Masking*: In some cases, it was useful to mask one organelle type with another. These masks could be inclusive (voxels outside the mask are set to background) or exclusive (voxels inside the mask are set to background). The masks could also be expanded by a set distance to increase the size of the mask. Masking proved especially useful when the optimal predictions for different classes came from different networks and iterations, meaning a single voxel could be assigned to multiple classes.

*Ribosomes*: For the purposes of analysis, ribosomes were treated as spheres with radius r = 10 nm. The first step in segmenting out the ribosomes was finding sphere centers in the predictions. To this end, we empirically derived a sphereness center metric (SCM) that seemed to work well locating likely ribosome centers. Obtaining the SCM was performed as follows:

We first thresholded the predictions at signed tanh distance transform ≥ 0. Let the set of all such above threshold voxels be called $V$. For each $v_i \in V$, we found all $v_{j\neq i} \in V$ that were both within $r$ = 10 nm of $v_i$and such that the shortest path from $v_i$ to $v_j$ did not cross through any background voxels. Let $n$ equal the number of such voxels satisfying this criterion for a given $v_i$, and $d_n$ be the distances between those voxels and $v_i$. The SCM was then calculated for each $v_i$ as follows:

$$SCM_{v_i} = n + \left(1 - \frac{1}{n}\sum_{d \epsilon d_n} \frac{d}{r}\right)$$

In this way, the sphereness center metric is heavily weighted by $n$ - the points in a ribosome-sized sphere that can be reached from $v_i$in a straight line through ribosome voxels - but it is also negatively weighted by $\frac{1}{n}\sum_{d\in d_n} \frac{d}{r}$- how far their center of mass is from the current voxel of interest, normalized by the radius $r$ = 10 nm.

Next, local maxima in SCM were found, with a requirement that no two local maxima could be closer than 12 nm of each other. The resulting points were considered ribosome centers and were expanded by 10 nm to create the spherical representation of the ribosomes.

*Microtubules:* Microtubule refinement as proposed in Eckstein et al. 2020 depends on solving a discrete optimization problem with the following associated hyperparameters: 1) A data dependent evidence prior *e*, 2) a stiffness factor *a*, controlling how much deviations from a straight line are punished, 3) A start edge prior *p*, defining the cost for starting and ending a trajectory, 4) a (negative) selection prior *c* needed to avoid the trivial solution of selecting no edges and no vertices 5) A data threshold *t* for candidate extraction and 6) a distance threshold *d* below which two candidates are connected by a candidate edge. For further details see Eckstein et al. 2020[28]. Hyperparameters used for each cross validation run presented in Extended Data Fig. 5, were found via grid search over two out of four densely traced 2 μm × 2 μm × 2 μm microtubule validation blocks in HeLa 2: *e* = 200, *a* = 22, *p* = 200 and *c* = -200, *t* = 0.4, d = 180 nm. The ILP was solved block wise with a size of 400 nm × 400 nm × 400 nm for each block.

In order to test applicability on our data, we compared Eckstein et al. to a baseline method consisting of: 1. Thresholding of the microtubule predictions, 2. Morphological closing of the thresholded predictions, 3. Connected component analysis, 4. Size filtering of connected components and 5. Skeletonization of each connected component, where we restrict the skeleton to have no branches. The baseline microtubule refinement used for comparison has hyperparameters: 1) Microtubule prediction threshold *t*, 2) Filter size *f* used for morphological closing of the thresholded predictions and 3) Size *c* for filtering of connected components. Grid search over the aforementioned validation blocks led to a majority winner setup with: *t* = 0.4, *c* = 4 and *f* = 500.

We evaluated the reconstructed microtubule tracks against the skeleton ground truth tracks by resampling both reconstruction and ground truth tracks equidistantly and subsequently matching vertices based on distance via Hungarian matching. Results were reported in terms of topological errors (Extended Data Fig. 6a), where errors were counted on the level of full microtubule trajectories, i.e. a trajectory was correct if it was matched to a reconstructed trajectory, and in terms of precision and recall on individual edges (Extended Data Fig. 6b), where an edge was correct, if the edge connected two matched vertices, that were matched to the same microtubule track. On average, the method reduced false positives, splits, and merges per micron of ground truth microtubule cable by a factor of ~2 w.r.t. to the baseline (Extended Data Fig. 6).

We expanded the microtubule tracks to create tubes with inner radius 6 nm and outer radius 12.5 nm, consistent with experimental measurements[73].

#### Quantifications

Unless otherwise specified, the following measurements were calculated for each unique organelle instance within a class.

*Instance count:* The number of unique instances of an organelle within a segmentation.

*Volume*: The total volume of all voxels within an organelle.

*Surface Area:* A given organelle's surface area is the surface area of all voxel faces that touch either the background or another organelle instance.

*Skeletons and Medial Surface:* Topological thinning to produce skeletons and medial surfaces was based on Lee et al.[47]. We used Skeletonize3D[46], the skeletonization implementation of Lee et al., as a starting point for our code. We updated it to remove bugs and added the ability to calculate medial surfaces. We also modified the implementation such that it could be run in parallel, based on Matlab Skeleton3D[74]. As input, we used a binarized version of the segmented volumes. In the event that two organelle instances touched, they were binarized and thinned independently. Topological thinning was applied to both mitochondria and ER (Extended Data Fig. 8).

*Length of Mitochondria*: Mitochondria skeleton lengths were analyzed using the Floyd Warshall algorithm. The skeletons were first cleaned by repeatedly pruning such that no remaining branch was shorter than 80 nm *(*Extended Data Fig. 8j,k). The mitochondrial length was then calculated as the longest shortest path within the pruned skeleton (Extended Data Fig. 8l).

*Diameter of Mitochondria:* To get mitochondria diameters, we first calculated the distance transform within the mitochondria. The radius is then the average distance transform along the longest shortest path within the pruned skeleton (see above), and the diameter is simply twice the radius.

*Morphology of ER*: To calculate the 3D membrane curvature of the ER (Extended Data Fig. 8e), we follow the methodology of Descoteaux et al.[36]. In brief, a planar metric is calculated for each voxel based on the Hessian matrix eigenvalues at that voxel. The metric was calculated over a scale space $\Sigma$ with a voxel's planarity taken to be its maximum planarity over the scale space:

$$planarity = max_{\sigma \epsilon \Sigma} \begin{cases} 0 & \lambda_3 > 0 \\ \exp(\frac{-R_{sheet}^2}{2\alpha^2})(1 - exp(\frac{-R_{blob}^2}{2\beta^2}))(1 - exp(\frac{-R_{noise}^2}{2c^2})) & \lambda_3 \leq 0 \end{cases}$$

, where $R_{plane} = \frac{|\lambda_2|}{|\lambda_3|}$, $R_{blob} = \frac{|(2|\lambda_3| - |\lambda_2| - |\lambda_1|)|}{|\lambda_3|}$, and $R_{noise} = \sqrt{\lambda_1^2 + \lambda_2^2 + \lambda_3^2}$ for eigenvalues$|\lambda_3| \geq |\lambda_2| \geq |\lambda_1|$. We set as constants$\alpha = 0.5, \beta = 0.5, c = 0.5$, used a binarized version of the segmented ER as the input image and smoothed over a scale space with 2 nm ≤ σ < 128 nm.

A consequence of this method was that planarity increases towards the center of planes[36]. However we wanted a planarity measure that is approximately constant for all voxels within a given planar (or non-planar) structure and is independent of distance from the center. To this end, we first calculated the medial surface of the ER via iterative thinning (see above) and restricted our planarity calculations to voxels on the medial surface (Extended Data Fig. 8f,g). Once planarity was calculated, we then created a reconstruction of the ER by expanding the medial surface by spheres centered at the medial surface with radii equal to the distance transform at each medial surface voxel. Each voxel within the reconstruction takes on the average planarity value of all spheres that contain it. An expert user then chose a threshold above which voxels were considered planes. Our threshold was 0.6 (Extended Data Fig. 8h).

To obtain a planarity measure of individual organelle or contact site instances, we took the mean planarity measure of all relevant voxels. For organelles, the relevant voxels were reconstructed ER voxels that are part of contact sites between the reconstructed ER and the given organelle instance. For contact sites, the relevant voxels were those contact site voxels that are part of the reconstructed ER.

When reporting planarity results in terms of surface area, we take the weighted average planarity of all relevant voxels, where the weights are the exposed surface area of each voxel.

Note: Volumes and surface areas of original and reconstructed ER differed by less than 10% and visual inspection indicated that the two matched well. Differences between the two are not expected to significantly affect the qualitative results.

*Contact Sites*: Generally a contact site is defined to be the region of space where two organelle instances (from different classes) are within a certain distance of each other. A naive approach to find contact sites would then be to define a contact site as a connected region of voxels that appear within the contact distance of both organelles, excluding all organelle voxels except those on the surface. An example of this implementation for contact sites between ER and mitochondria is shown in Extended Data Fig. 8a-d. This approach has two major drawbacks. First, a voxel may be considered part of a contact site when it is within the contact distance of both organelles even if the organelles themselves are separated by more than the contact distance. For example, if two organelle voxels are 15 nm apart and a contact distance of 10 nm is chosen, some voxels will be within 10 nm of both organelles even though the organelles themselves are 15 nm apart. This can lead to floating contact sites that aren't touching any organelle as well as overly extended contact sites. The second disadvantage of this method is that it does not restrict contact sites to exist only where organelle surfaces are facing each other. That is, a voxel may be considered part of a contact site so long as it is within the contact distance even if it is on the far side of the organelle without a direct line of sight to the contacting organelle.

We solved these issues with a different approach for measuring contact sites. First, we find all organelle surface voxels in one class (A) that are within the cutoff distance of organelles in the other class (B). These were considered contact site surface voxels (CSSV). For each contacting A-B pair, we created a binary image of their CSSV. We then filled in the remaining contact site voxels by connecting the corresponding CSSV from A to B in the binary image using an extension of Bressenham's line algorithm[75] to 3D. For a given pair of surface voxels, we only kept the line if 1) the distance between the surface voxels was less than or equal to the cutoff distance and 2) the line did not cross through any organelle voxels (except the start and end surface voxels). Combined, these two criteria fixed the issues mentioned above and the results can be seen in Extended Data Fig. 8d. For simplicity, contact-site surface voxels in one class are only assumed to be in contact with the nearest neighboring organelle of the other class.

Since our approach involved lines and surfaces but takes place in a discrete voxel space, it is not perfect; for example, it can result in contact sites that contain holes. Nevertheless, for our purposes, it was a more appropriate measurement than that produced with the naive approach.

Note that when calculating distances involving surface voxels, the contact distance was extended by 4 nm to account for the thickness of the surface voxels themselves. Additionally, organelles are often pulled from different networks/iterations meaning they can occupy the same space. Thus in many cases, we mask out the thinner organelle with the thicker one in order to prevent complete removal of contact sites in the event of overlap.

#### CLEM Registration

Automatic registration of light (LM) and electron microscopic (EM) images is challenging due to resolution and contrast differences between the modalities, and non-linear transformations induced during sample preparation. We generated synthetic images derived from the EM segmentation that resemble light images of fluorescent organelle markers, suitable for automatic voxel-based deformable registration algorithms. In this work, image data came from a previously published COS7 cell transiently expressing ER luminal and mitochondria membrane markers, mEmerald-ER3 and Halo/JF525-TOMM20 respectively[39]. We registered the mitochondria membrane class in the prediction to the fluorescent channel labeling TOMM20. The LM image was the algorithm's registration target. This served to correct any shape or morphology changes that occur during the preparation for electron microscopy, since the LM images are more likely to represent the true morphology than the EM due to more "gentle" preparation.

Rather than using EM mitochondria predictions directly, we used them to generate "synthetic LM" images which improve the downstream registration algorithm's efficiency and performance. We first downsampled the prediction from the EM to a resolution close to that of the light (64 nm x 64 nm x 64 nm) and reoriented it so its coordinate directions match that of the light image. Next, we applied an anisotropic Gaussian blur (kernel width 1 x 1 x 3 voxels at the downsampled resolution) filter to the downsampled predictions to mimic the point spread function of the light microscope. These steps were performed using Fiji[53].

The EM and LM images used in this study contained several cells in the field of view, but one cell was not fully labeled by the fluorescent marker and was therefore "invisible" in the LM image. The learned network could successfully segment mitochondria membrane from the EM, meaning the EM and LM images consisted of different image content, posing a significant challenge for image registration. We manually masked (removed) those parts of the EM segmentation that have no corresponding signal in the LM using Fiji[53]. This intervention cost less than thirty minutes of human attention and improved the automatic registration that follows.

We used elastix[48], a state-of-the-art, automated registration algorithm, with the synthetic image derived from mitochondria as the moving image, and the manually masked LM TOMM20 image as the target. We ran elastix in two steps, where the first estimates an affine transformation. The second step was initialized with the affine and finds a non-linear transformation mapping the moving EM image to the target LM image. We used normalized cross correlation as the image similarity metric. Specific parameter configurations can be found online (https://github.com/janelia-cosem/cosem-lm-em-registration). Extended Data Fig. 9 shows the results of this automatic alignment.

The computational expense was larger for transforming the full resolution EM images and predictions and was not possible using built-in elastix tools due to their size. Therefore, we wrote software for transforming very large image data that use the N5 block-based file format[42], and parallelizes over blocks using Apache Spark[76]. We also used N5 to store the transformation itself as a displacement field[77] because it enables the loading of only the small subset of a transformation needed for each block, substantially reducing overhead.

The Jacobian determinant is a commonly used measure of image deformation. The Jacobian determinant for the CLEM spatial transformation is visualized in Fig. 4e,f. Notice that it takes values between about 0.6 and 1.4, indicating that space is locally stretched or shrunk by up to 40%, though the vast majority lie between 0.8 and 1.2. The small variance of this distribution (0.003) indicates that the transformation is smooth overall. For registration algorithms that are capable of producing large distortions (as ours is), smooth results are an indicator of promising results.

We measured accuracy by comparing the automatically generated transformation with human generated ground truth. One human annotator manually placed 31 landmarks on the ER light images, and 31 points at the corresponding locations on the ER predictions generated from EM images. The landmark locations for the light image were given to a second human rater, who manually found the corresponding points on the EM-derived ER prediction image. This gives us two independent estimates of the transformation at those 31 selected locations. Applying the automatically generated transformation to those 31 points and measuring the distance of the result to the ground-truth (human) corresponding point gives an estimate of the accuracy of the automatic transformation.

Fig. 4g shows the error magnitudes of the automatic registration at those landmarks and the inter-annotator errors. For these points, the errors made by the automatic registration were larger than the inter-annotator errors, in part because landmark locations were selected specifically so that they could be localized consistently. Those locations were mostly at junctions of the ER network.

Notice also that human annotators placed landmarks using ER images whereas the automatic algorithm had access only to mitochondria images. This makes the human ground-truth transformation independent of the automatically generated transform. The resulting error estimate therefore reflects both the algorithm's ability to register its input images of mitochondria, but also the extent to which those images are indicative of the true transformation over the field of view of the image. For example, the right half of the field of view of Fig. 4g lacks mitochondria signal, and unsurprisingly shows (qualitatively) higher errors. We expect registration to be accurate only where there exists a signal from which to estimate a transformation.

Two light microscopic imaging modalities (PALM and SIM) were collected for this sample, enabling us to measure the consistency of our automated registration procedure for changes in the LM imaging modality. We independently registered the "synthetic" image derived from mitochondria predictions to each of the two light modalities and computed spatial error . The spatial error at each point is the distance between the result of transforming that point with each of the two independent transformations. Fig. 4h shows a spatial map of these errors. Fig. 4i shows a histogram of these errors inside the cell. The mode of the error distribution is less than

0.2 μm, indicating that the registration procedure was consistent across the two light modalities. Regions with larger errors approaching 0.6 μm occur (rarely) near the periphery of the cell, far from the mitochondria whose signal drives the registration. The larger errors of about 1.0 μm occurring outside of the cells are of no concern because they cannot affect any biologically relevant analyses.

A single choice for registration parameters did not perform equally well for both LM modalities tested (PALM and SIM). This was to be expected because these modalities differ in extent and kind of imaging artifacts. Finding a good set of parameters to accurately register a given pair of images is often difficult and time consuming. The difficulty lies in the expertise needed to improve parameters in response to a particular kind of error. For example, what parameters need to change if the transformation applies too large a distortion, or is not sufficiently flexible? An automated grid search is possible only if a good and objective metric is available to assess the result, but such a metric is not usually available. To expedite this process, we created a varied set of registration parameters and a set of scripts that enables a human to run a registration task with a large set of parameters, and to quickly view all the results at once. This minimizes human parameter adjustment at the cost of computation time. The additional computational cost is small: elastix completed in less than five minutes using a typical set of parameters for images of the size and resolution used here.

### Supplementary Discussion

#### Training data

Our experiments show that this work is promising but not finished. While our networks perform excellently on datasets and structures that are well represented in the training data, performance degrades for rare structures and in samples whose properties are not well represented (see Fig. 2a,b). This was expected and can be addressed by surgically adding more training data for atypical samples and rare structures such that, over time, everything that is structurally possible will be reasonably well represented. Over the course of our experiments, we have seen consistent improvement of our networks' performance as we added new training examples which indicates that adding more examples will remain helpful to improve generalization.

How much will be enough? Today, our public training data comprises 73 annotated volumes sampled from 5 different cell types, overall ~635 Mvx, each annotated with 35(37) class labels. At first glance, this seems to compare favorably to e.g. the the largest publicly available training volume for neuron segmentation and synapse detection from EM (CREMI) with ~586 Mvx.[60] However, the CREMI training data annotates only three categories (cell boundaries, synaptic clefts, and synaptic partners), and is dedicated to just one, albeit very large EM sample. The largest ground truth volume capturing the variability of mitochondria in neural tissue is the recently published ~28 Gvx large MitoEM dataset.[19] This dataset focuses on two specific neural tissue samples and annotates mitochondria instances, not their membranes or other organelles. The authors observed an astonishing variability in shape and appearance of mitochondria in these samples, justifying at least some of the large sample size. In this light, is it realistic to expect the OpenOrganelle training data to reach a volume that will be sufficient to represent the impressive variability of cellular structures across cell types and tissues? Will our networks be able to learn this? We are optimistic. So far, adding more training data has strictly improved performance. We are therefore working on expanding the dataset to tissue samples, and we continuously add annotations on samples and organelles that currently perform unsatisfactorily. However, we believe that this will remain an ongoing effort for some time to come and that both improved data and innovative new methods will contribute to solving this important problem. Just as ImageNet[78] has revolutionized computer vision research by being representative for everyday object classification from images, we expect OpenOrganelle to become a comprehensive and representative dataset for all cellular organelles that can be reconstructed from high resolution FIB-SEM of cells and tissues, and we invite the scientific community to contribute data, annotations, and methods to this ongoing effort.

#### Augmentation

It is well understood that appropriate augmentation of training data helps to make networks become invariant or robust with respect to the augmented properties. However, it is often unclear what appropriate augmentations are in a concrete application. In the experiments described here, we have focused on intensity variations, geometric distortions, and, in the case of the 8 nm training, on some amount of imaging noise. It is clear that we have to do more. Preliminary ongoing experiments suggest that noise free data generally leads to better performance, and we plan to use state-of-the-art denoising methods[79,80] trained directly on noisy 3D data to preprocess our volumes. While noise-augmentation on the training data should theoretically achieve the same result, de-noising, if treated independently, can be trained on annotation free samples from the entire dataset which dramatically increases the training volume.

Generally, better input data leads to better reconstructions. Besides noise, some of our volumes suffer from inconsistent contrast, imperfect alignment, and compression artifacts from variation in how FIB milling progresses. The quality of organelle reconstruction strongly depends on the presence of these artifacts. We will therefore spend additional efforts on improving input data quality, applying and expanding upon state-of-the-art methods for contrast correction, alignment, and compression correction.

#### Balancing, network architectures, and validation

Balancing training data matters and is difficult. While the frequency of individual structures and shapes can be an important property for our networks to learn, it is also important to make sure that, during the course of training, all classes are shown often enough such that they can be ‘understood’ and will not be ‘forgotten’ in favor of others. It is difficult to achieve balance in a training setup with many classes that occur at vastly different frequencies (see Extended Data Fig. 2a). Our current training pipeline for multi-class networks does not include explicit methods for balancing relative class frequency resulting in rare classes being less well reconstructed than frequent classes. The frequency of positive and negative labels of each individual class is normalized per training patch, saturating at 5% and 95%, respectively. Three of our few-class networks were trained with a balanced frequency of drawing samples from training blocks that contain positive annotations and those that do not. Without this importance sampling, the networks did not succeed to learn the task which underlines the importance of presenting rare classes at increased frequency.

Meaningful balancing is further complicated by the fact that we train our networks to predict tanh signed boundary distances instead of voxel independent class probabilities. While this has the advantage to promote meaningful object shapes, and trains faster[10], we should balance the frequency of all possible distance predictions instead of class frequency. Class frequency is only a poor surrogate for signed boundary distances that capture shape properties such as the average proximity of object instances, their size, or compactness. The downside of this expressive power is that tanh signed boundary distances are not invariant with respect to geometric distortion and therefore cannot be precomputed for the entire dataset. Calculating the training signal for all 35 classes is computationally demanding and currently requires that several CPU workers support a single GPU during training. We can therefore not afford to reject training patches based on their cumulative properties. As a workaround, we are currently working on generating approximate precomputed sample probability maps over all training data that we plan to use for importance sampling considering all classes and distances.

One curious observation was that the performance of our networks on 8 nm data is typically as good as or, for some organelles, better than on 4 nm data. This suggests that the FOV of the network can be more important than high resolution detail. We are currently working on architectures that increase the FOV without increased demands on GPU memory to exploit this. We also try to find the 'optimal' resolution for each organelle, e.g. identifying and reconstructing large organelles such as mitochondria is likely to require significantly lower resolution than identifying and reconstructing microtubules, ribosomes, or nuclear pores.

Our manual validation procedure sampling proofreader sentiment to compare training iterations and architectures has proven to be an efficient alternative to validation on dedicated ground truth data. This is particularly important to assess the performance of an established network when applied to previously unseen data where such ground truth will typically not be available. In our experiments, manual and ground truth based validation are generally in agreement, however, it is important to understand that we do not yet know when it saturates. It is relatively easy to rate a bad performing and a well performing network, but it becomes increasingly difficult to make a decision about two very well performing networks. On the other hand, the manual validation procedure samples from the entire dataset, which avoids a clear shortcoming of our ground truth based validation: The holdout blocks, at times, fall short to capture intra-dataset variability and a sufficient number of variable organelle instances. This is particularly visible for mitochondria in *jrc_macrophage-2* (see Fig 2a, Extended Data Fig. 3a) where the holdout block contains only part of an atypical mitochondria. It is also important to understand that the metric used for ground truth based validation can have an equally unpredictable impact as we showed for F1 score and Mean false distance (see Supplementary Methods: Evaluations, Extended Data Fig. 3b). Overall, the combination of sentiment based manual validation and follow-up ground truth based validation has proven to be a powerful tool to develop and improve our training setups.

### Supplementary Videos

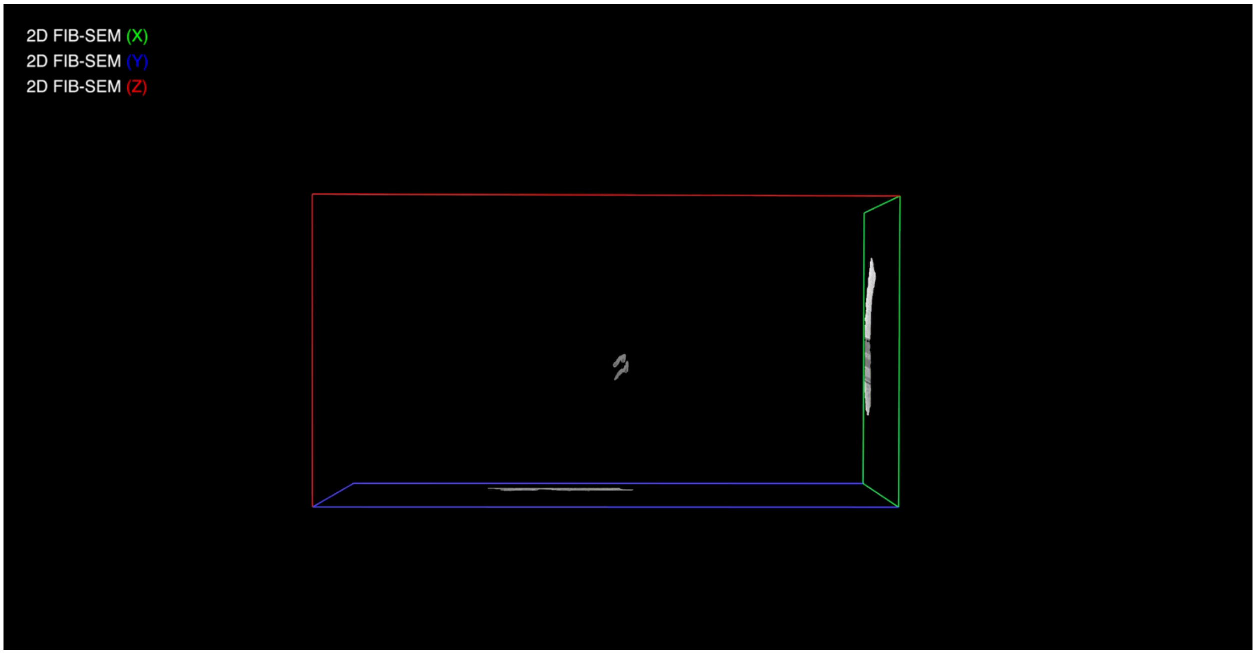


#### Supplementary Video 1 - FIB-SEM, training data, and predictions

Showcase of the processing pipeline, using *jrc_hela-2* as an example. We begin with the EM data. Then skilled annotators carefully classify every voxel within a volume; shown here are 15 of these training blocks. These segmentations are fed into machine learning algorithms as training data. The prediction output from these algorithms are refined. Once the predicted, whole-cell segmentations are achieved, quantitative analytics of subcellular distributions, interactions, sizes, and morphologies can be acquired as shown in Supplementary Video 2.

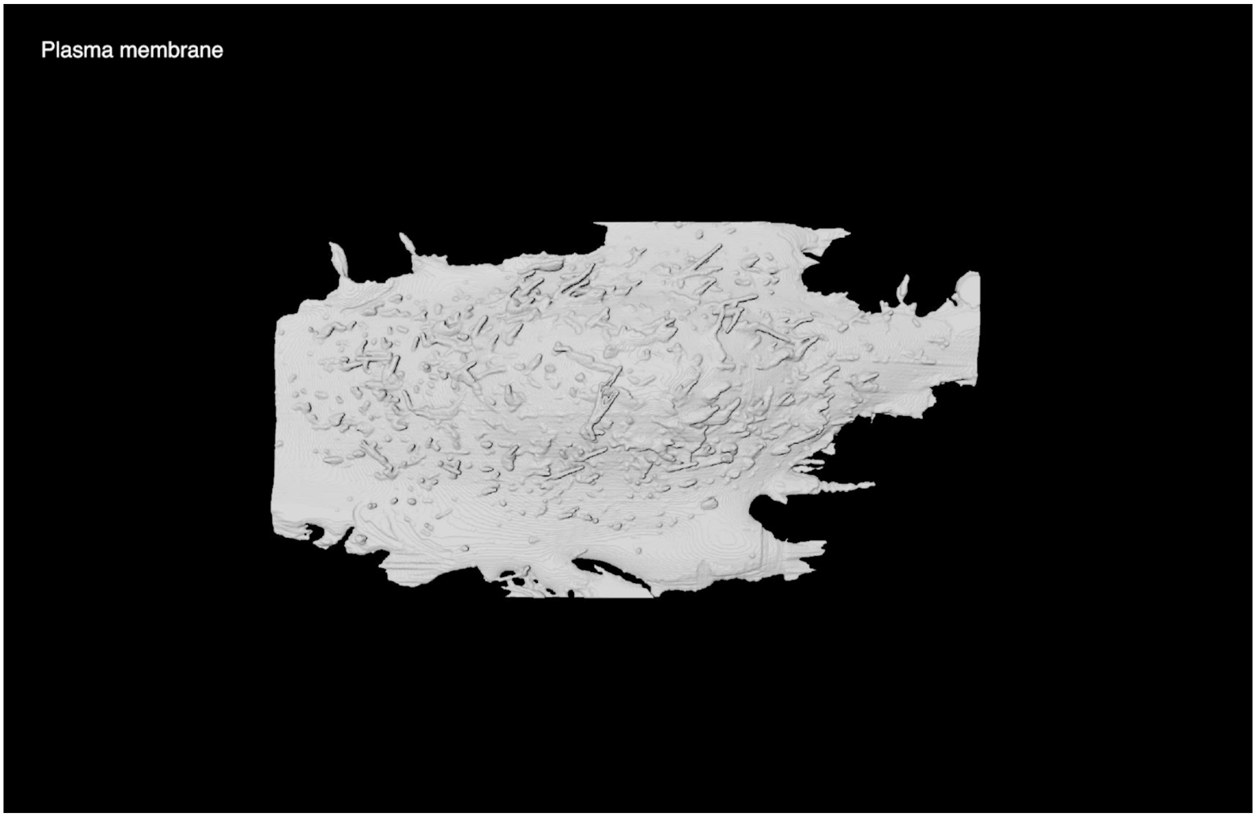


#### Supplementary Video 2 - Three analysis examples

Three analysis examples in *jrc_hela-2*. The first example is from Fig. 3b, a microtubule contacting multiple different organelles. The second example is from Fig. 3d, displaying the relationship between ER morphology and mitochondria contact sites. The third example is from Fig. 3h, showing the distribution of ribosomes bound to the ER.

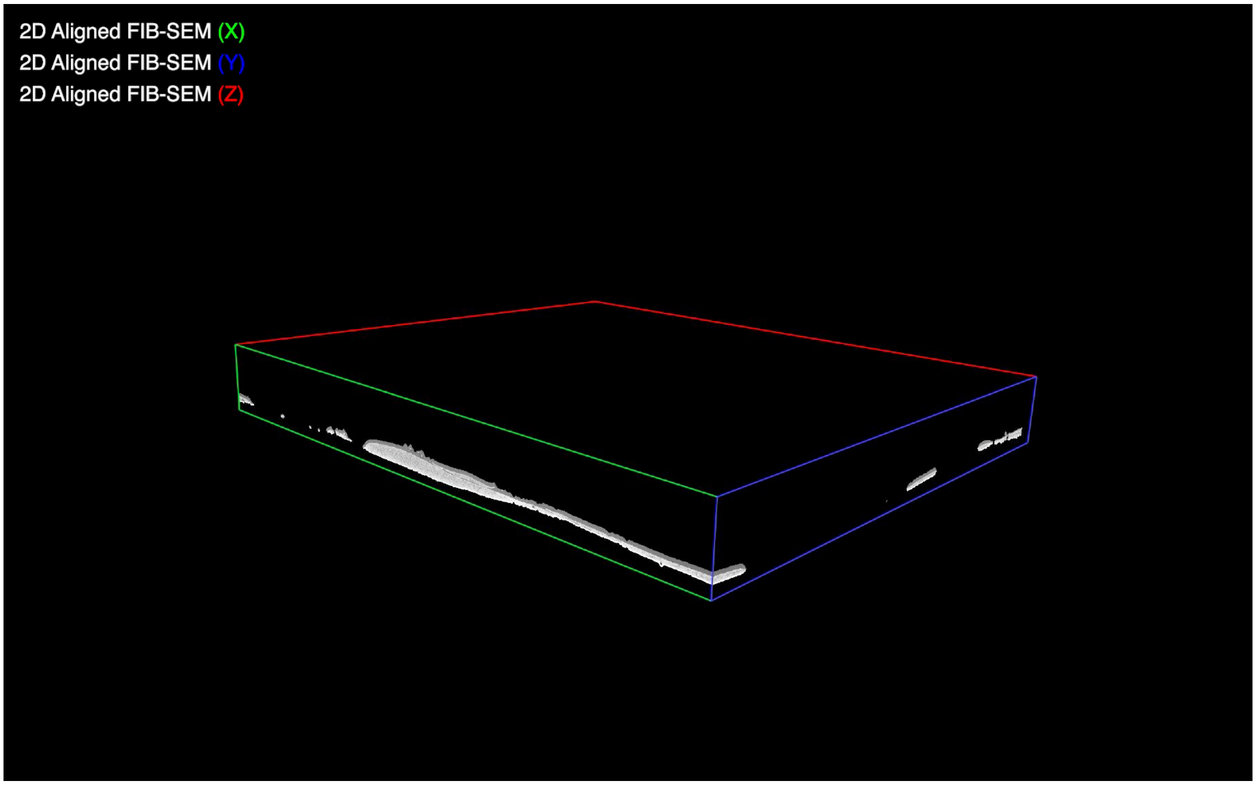


#### Supplementary Video 3 - CLEM registration

FIB-SEM and correlative light microscopy automatically registered using whole cell mitochondria membrane predictions. Displayed are PALM images of mitochondria membrane marker and Halo/JF525-TOMM20 and ER luminal marker mEmerald-ER3, predictions for mitochondria membrane and ER, as well as the corresponding (8 nm × 8 nm × 8 nm) FIB-SEM. A "warping" from affine-only to the full-deformable transformation is also shown.

### Supplementary Tables

| Class | All (A) | Many (M) | Few (F) | | | | | | | | | | | | | | Best network and iteration | | | | | | |
|---|---|---|---|---|---|---|---|---|---|---|---|---|---|---|---|---|---|---|---|---|---|---|---|
| | | | | | | | | | | | | | | | | | HeLa 2 | HeLa 3 | Jurkat | Mac | HeLa1 | Ch. Pl. | COS-7 |
| Centrosome | • | | • | | | | | | | | | | | | | | A975k | A600k | A600k | A600k | – | – | – |
| Centrosome D App | • | | • | | | | | | | | | | | | | | A900k | A900k | A900k | A900k | – | – | – |
| Centrosome SD App | • | | • | | | | | | | | | | | | | | A1075k | / | / | / | – | – | – |
| Chromatin | • | | | • | | | | | | | | | | | | | A600k | A825k | A825k | A825k | – | – | – |
| ECS | • | • | | | • | | | | | | | | | | | | M625k | F725k | M875k | A725k | – | – | – |
| Endo | • | • | | | | • | | | | | | | | | | | M925k | M925k | M650k | M650k | M975k | M625k | – |
| Endo mem | • | • | | | | • | | | | | | | | | | | M1025k | M1025k | M1025k | M1025k | – | – | – |
| ER | • | • | | | | | • | | | | | | | | | | M625k | A1075k | F625k | M650k | M925k | M650k | F825k |
| ER mem | • | • | | | | | • | | | | | | | | | | M625k | A625k | A775k | A775k | – | – | – |
| ERES | • | • | | | | | | • | | | | | | | | | A925k | M750k | A875k | A850k | – | – | – |
| ERES mem | | | | | | | | • | | | | | | | | | / | / | / | / | – | – | – |
| E Chrom | • | | | • | | | | | | | | | | | | | / | / | / | / | – | – | – |
| N-E Chrom | • | | | • | | | | | | | | | | | | | / | / | / | / | – | – | – |
| Golgi | • | | | | | | | | • | | | | | | | | F650k | F650k | F650k | F650k | – | – | – |
| Golgi mem | • | | | | | | | | • | | | | | | | | F650k | F650k | F650k | F650k | – | – | – |
| H Chrom | • | | | | | | | | | | | | | | | | / | / | / | / | – | – | – |
| N-H Chrom | • | | | • | | | | | | | | | | | | | A1200k | A600k | A600k | A600k | – | – | – |
| LD | • | | | | | | | | | • | | | | | | | A1175k | A925k | A525k | A925k | – | – | – |
| LD mem | • | | | | | | | | | • | | | | | | | A625k | A625k | A625k | A875k | – | – | – |
| Lyso | • | | | | | • | | | | | | | | | | | A900k | A1125k | A1125k | A725k | – | – | – |
| Lyso mem | • | | | | | • | | | | | | | | | | | A900k | A700k | A1125k | A1125k | – | – | – |
| MT | • | • | | | | | | | | | • | | | | | | F1400k | F1300k | F900k | A1100k | – | – | – |
| MT out | • | • | | | | | | | | | • | | | | | | F1325k | F1300k | F975k | / | – | – | – |
| Mito | • | • | | | | | | | | | | • | | | | | F575k | A825k | A875k | M1100k* | M875k | M875k | F750k |
| Mito mem | • | • | | | | | | | | | | • | | | | | A900k | A925k | A925k | M650k | – | – | F650k |
| Mito Ribo | • | | | | | | | | | | | • | | | | | / | / | / | / | – | – | – |
| NE | • | | | | | | | | | | | | • | | | | A750k | A1150k | A1150k | A725k | – | – | – |
| NE mem | • | | | | | | | | | | | | • | | | | A1150k | A1150k | A1150k | A1150k | – | – | – |
| NP | • | | | | | | | | | | | | • | | | | A1075k | A1075k | A900k | A900k | – | – | – |
| NP out | • | | | | | | | | | | | | • | | | | A975k | A975k | A975k | A975k | – | – | – |
| Nucleolus | • | | | | | | | | | | | | | • | | | A950k | A600k | A950k | F625k | – | – | – |
| Nucleus | • | • | | | | | | | | | | | | • | | | A600k | M625k | F600k* | F650k* | M575k | M1100k | – |
| PM | • | • | | | • | | | | | | | | | | | | M650k | M650k | M650k | M650k | M975k | M975k | – |
| Ribo | • | | | | | | | | | | | | | | • | | F525k | A1200k | F575k | F525k | – | – | – |
| Vesicle | • | • | | | | | | | | | | | | | | • | F675k | F675k | F675k | F550k | M825k | M800k | – |
| Vesicle mem | • | • | | | | | | | | | | | | | | • | A1075k | F575k | A1075k | A1075k | – | – | – |

#### Supplementary Table 1 - Best network and iteration per class and dataset

Classes included in each trained network and best network and iteration for each class and dataset chosen via manual evaluation. Predictions that deviated from the manual evaluation are denoted by an (*). These predictions were re-evaluated to bias towards errors that our refinement pipeline could mitigate. (-) denotes setups that have not been evaluated, (/) denotes setups where no predictions were above threshold. The (8 nm × 8 nm × 8 nm) datasets were only evaluated for the “many” class network.

| | F1 Score | | | | Precision | | | | Recall | | | | Mean False Distance (nm) | | | |
|---|---|---|---|---|---|---|---|---|---|---|---|---|---|---|---|---|
| | *jrc_hela-2* | *jrc_hela-3* | *jrc_jurkat-1* | *jrc_macrophage-2* | *jrc_hela-2* | *jrc_hela-3* | *jrc_jurkat-1* | *jrc_macrophage-2* | *jrc_hela-2* | *jrc_hela-3* | *jrc_jurkat-1* | *jrc_macrophage-2* | *jrc_hela-2* | *jrc_hela-3* | *jrc_jurkat-1* | *jrc_macrophage-2* |
| *Chromatin* | 0.71 | 0.42 | 0.22 | 0.22 | 0.64 | 0.42 | 0.91 | 0.81 | 0.79 | 0.42 | 0.12 | 0.13 | 9.4 | 12 | 16 | 38 |
| *ECS* | 0.98 | 0.98 | 0.98 | 0.97 | 1.00 | 0.96 | 0.96 | 1.00 | 0.96 | 1.00 | 0.99 | 0.94 | 0.14 | 0.16 | 0.8 | 0.5 |
| *Endo* | 0.52 | 0.019 | 0.90 | 0.21 | 0.51 | 0.01 | 0.87 | 0.18 | 0.54 | 0.07 | 0.95 | 0.25 | 100 | 580 | 5.3 | 260 |
| *Endo mem* | 0.34 | 8.7E-3 | 0.53 | 0 | 0.67 | 0.011 | 0.72 | 0 | 0.23 | 7.3E-3 | 0.42 | 0 | 44 | 550 | 12 | 750 |
| *ER* | 0.84 | 0.71 | 0.75 | 0.97 | 0.79 | 0.90 | 0.79 | 0.87 | 0.90 | 0.58 | 0.71 | 0.87 | 2.5 | 8.8 | 25 | 3.4 |
| *ER mem* | 0.34 | 0.19 | 0.22 | 0.011 | 0.61 | 0.76 | 0.56 | 0.83 | 0.24 | 0.11 | 0.13 | 5.6E-3 | 8.3 | 19 | 37 | 24 |
| *ERES* | 0.33 | 0.12 | - | 0 | 0.45 | 0.17 | - | 0 | 0.25 | 0.095 | - | 0 | 230 | 170 | - | NaN |
| *Lyso* | 0.3 | 0.92 | 0.37 | 0.81 | 0.25 | 0.91 | 0.54 | 0.78 | 0.39 | 0.93 | 0.28 | 0.84 | 95 | 20 | 31 | 27 |
| *Lyso mem* | 0.21 | 9.3E-3 | 0.03 | 0.044 | 0.60 | 0.48 | 0.42 | 0.46 | 0.13 | 4.7E-3 | 0.015 | 0.023 | 93 | 59 | 45 | 54 |
| *Mito* | 0.93 | 0.97 | 0.98 | 0 | 0.95 | 0.97 | 0.99 | 0 | 0.91 | 0.96 | 0.96 | 0 | 3.6 | 0.31 | 0.15 | NaN |
| *Mito mem* | 0.64 | 0.67 | 0.60 | 0 | 0.76 | 0.72 | 0.78 | 0 | 0.55 | 0.63 | 0.49 | 0 | 14 | 3.7 | 6.8 | NaN |
| *NE* | 0.85 | 0.61 | 0.74 | 0.8 | 0.74 | 0.75 | 0.62 | 0.81 | 0.98 | 0.51 | 0.93 | 0.78 | 0.95 | 10 | 3 | 2.3 |
| *NE mem* | 0.68 | 0.27 | 0.51 | 0.18 | 0.62 | 0.39 | 0.41 | 0.54 | 0.76 | 0.20 | 0.67 | 0.11 | 1.8 | 17 | 4.3 | 16 |
| *NP* | 0.35 | 0.018 | 0.23 | 0.46 | 0.22 | 0.13 | 0.13 | 0.45 | 0.85 | 9.9E-3 | 1.00 | 0.46 | 21 | 290 | 25 | 24 |
| *NP out* | 0 | 0 | 0 | 0 | 0 | 0 | 0 | 0 | 0 | 0 | 0 | 0 | NaN | NaN | NaN | NaN |
| *Nucleus* | 0.99 | 0.98 | 0.91 | 0.95 | 0.98 | 0.97 | 0.97 | 0.92 | 1.00 | 0.99 | 0.85 | 0.99 | 0.095 | 0.13 | 1.5 | 0.74 |
| *PM* | 0.76 | 0.88 | 0.80 | 0.93 | 0.71 | 0.96 | 0.76 | 0.90 | 0.83 | 0.81 | 0.86 | 0.96 | 1.2 | 0.83 | 1.5 | 0.95 |
| *Ribo* | 0.3 | 0.25 | 0.23 | 0.24 | 0.20 | 0.19 | 0.19 | 0.17 | 0.56 | 0.39 | 0.29 | 0.38 | 11 | 15 | 17 | 12 |
| *Vesicle* | 0.12 | 0.31 | 0.07 | 0.13 | 0.50 | 0.45 | 0.15 | 0.32 | 0.066 | 0.24 | 0.045 | 0.08 | 170 | 150 | 280 | 169 |
| *Vesicle mem* | 0.2 | 0.095 | 9.9E-4 | 0 | 0.49 | 0.43 | 0.18 | 0 | 0.12 | 0.054 | 5.0E-4 | 0 | 73 | 140 | 220 | NaN |

#### Supplementary Table 2 - Evaluation metrics

F1 Score, Precision, Recall and Mean False Distance on holdout blocks from four datasets after refinements described in Supplementary Methods: Refinements. Network type and iterations represented here are listed in Supplementary Table 1 and were optimized manually with a bias towards potential improvement through the refinement process. See Extended Data Fig. 6 and Supplementary Table 4 for MTs.

| Dataset | Refinement | Centrosome | Centrosome D App | Chromatin | N-H Chrom | ECS | Endo | Endo mem | ER | ER mem | ERES | Golgi | Golgi mem | LD | LD mem | Lyso | Lyso mem | Mito | Mito mem | MT | NE | NE mem | NP | Nucleus | PM | Ribo | Vesicle | Vesicle mem |
|---|---|---|---|---|---|---|---|---|---|---|---|---|---|---|---|---|---|---|---|---|---|---|---|---|---|---|---|---|
| *jrc_hela-2* | Smoothing | | | • | | | • | | | | | | | • | | • | | • | | | | | | • | | | | |
| | Connected Components | • | • | • | • | • | • | | • | | • | • | | • | | • | | | | | • | | • | • | • | • | • | |
| | Size Filtering | • | • | • | • | • | • | | • | | • | • | | • | | • | | • | | | • | | • | • | • | | • | |
| | Hole Filling | | | | | | | | | | | | | | | | | • | | | | | | | | | | |
| | Custom Filtering | | | | | | | | | | | | | | | | | | | | | | | | • | | | |
| | Watershed and Agglomeration | | | | | | | | | | | | | | | | | • | | | | | | | | | | |
| | Masking | | | • | • | | • | | | | • | | | | | | | | | | • | | • | | | | | |
| | Masked to Parent Organelle | | | | | | | • | | • | | | • | | • | | • | | • | | | • | | | | | | • |
| | Custom Reconstruction | | | | | | | | | | | | | | | | | | | ** | | | | | | * | | |
| *jrc_hela-3* | Smoothing | | | • | | | • | | | | | | | • | | • | | • | | | | | | • | | | | |
| | Connected Components | • | • | • | • | • | • | | • | | • | • | | • | | • | | | | | • | | • | • | • | • | • | |
| | Size Filtering | • | • | • | • | • | • | | • | | • | • | | • | | • | | • | | | • | | • | • | • | | • | |
| | Hole Filling | | | | | | | | | | | | | | | | | • | | | | | | | | | | |
| | Custom Filtering | | | | | | | | | | | | | | | | | | | | | | | | • | | | |
| | Watershed and Agglomeration | | | | | | | | | | | | | | | | | • | | | | | | | | | | |
| | Masking | | | • | • | | • | | | | • | | | | | | | | | | • | | • | | | | | |
| | Masked to Parent Organelle | | | | | | | • | | • | | | • | | • | | • | | • | | | • | | | | | | • |
| | Custom Reconstruction | | | | | | | | | | | | | | | | | | | | | | | | | * | | |
| *jrc_jurkat-1* | Smoothing | | | • | | | • | | | | | | | • | | • | | • | | | | | | • | | | | |
| | Connected Components | • | • | • | • | • | • | | • | | • | • | | • | | • | | | | | • | | • | • | • | • | • | |
| | Size Filtering | • | • | • | • | • | • | | • | | • | • | | • | | • | | • | | | • | | • | | • | | • | |
| | Hole Filling | | | | | | | | | | | | | | | | | • | | | | | | | | | | |
| | Custom Filtering | | | | | | | | | | | | | | | | | | | | | | | • | • | | | |
| | Watershed and Agglomeration | | | | | | | | | | | | | | | | | • | | | | | | • | | | | |
| | Masking | | | • | • | | • | | | | • | | | | | | | | | | • | | • | | | | | |
| | Masked to Parent Organelle | | | | | | | • | | • | | | • | | • | | • | | • | | | • | | | | | | • |
| | Custom Reconstruction | | | | | | | | | | | | | | | | | | | | | | | *** | | * | | |
| *jrc_macrophage-2* | Smoothing | | | • | | | • | | | | | | | • | | • | | • | | | | | | • | | | | |
| | Connected Components | • | • | • | • | • | • | | • | | • | • | | • | | • | | | | | • | | • | • | • | • | • | |
| | Size Filtering | • | • | • | • | • | • | | • | | • | • | | • | | • | | • | | | • | | • | • | • | | • | |
| | Hole Filling | | | | | | | | | | | | | | | | | • | | | | | | | | | | |
| | Custom Filtering | | | | | | | | | | | | | | | | | | | | | | | | • | | | |
| | Watershed and Agglomeration | | | | | | | | | | | | | | | | | • | | | | | | | | | | |
| | Masking | | | • | • | | • | | | | • | | | | | | | | | | • | | • | | | | | |
| | Masked to Parent Organelle | | | | | | | • | | • | | | • | | • | | • | | • | | | • | | | | | | • |
| | Custom Reconstruction | | | | | | | | | | | | | | | | | | | | | | | | | * | | |

#### Supplementary Table 3- Segmentation refinements used

List of refinements performed for organelles from four datasets.
*See Supplementary Methods: Refinements - Ribosomes.
**See Supplementary Methods: Refinements - Microtubules.
***After watershedding and agglomeration, an expert-user selected large objects to keep. These objects, and any objects touching them, were kept as this proved ideal for this nucleus.

| | | Precision | Recall | F1 | FPs/Length ($\mu m^{-1}$) | FNs/Length ($\mu m^{-1}$) | Splits/Length ($\mu m^{-1}$) | Merges/Length ($\mu m^{-1}$) |
|---|---|---|---|---|---|---|---|---|
| *jrc_hela-2* | Region 1 | 0.94 | 0.47 | 0.62 | 0.12 | 0.56 | 0.59 | 0.26 |
| | Region 2 | 0.95 | 0.71 | 0.81 | 0.11 | 0.55 | 0.98 | 0.49 |
| | Region 3 | 0.85 | 0.19 | 0.31 | 0.05 | 0.71 | 0.25 | 0.11 |
| | Region 4 | 0.91 | 0.46 | 0.61 | 0.12 | 0.47 | 0.41 | 98 |
| *jrc_hela-3* | Region 1 | 0.91 | 0.53 | 0.67 | 0.13 | 0.52 | 0.65 | 65 |
| | Region 2 | 0.89 | 0.71 | 0.79 | 0.17 | 0.37 | 0.55 | 0.2 |
| *jrc_jurkat-1* | Region 1 | 0.99 | 0.24 | 0.39 | 0.03 | 0.96 | 0.29 | 0.07 |
| | Region 2 | 0 | 0 | 0 | 0 | 0.96 | 0 | 0 |
| *jrc_macrophage-2* | Region 1 | 0.65 | 0.23 | 0.34 | 0.35 | 0.89 | 0.52 | 0.22 |
| | Region 2 | 0 | 0 | 0 | 0 | 1.2 | 0 | 0 |

#### Supplementary Table 4 - Microtubule evaluation metrics

Precision, Recall, F1 Score, and topological errors normalized by ground-truth microtubule cable length for each cell.

| Dataset | Measurement | *ER* | *Golgi* | *Mito* | *Nucleus* | *PM* |
|---|---|---|---|---|---|---|
| *jrc_hela-2* | Total Volume (µm³) | 130 | 5.8 | 110 | 610 | 81 |
| | Total Surface Area (µm²) | 1.0E4 | 490 | 1.9E3 | 850 | 1.0E4 |
| | Number of Objects | 24 | 8 | 421 | 3 | 3 |
| | Mean Volume ± SD Per Object (µm³) | 5.3 ± 21 | 0.72 ± 1.7 | 0.26 ± 0.48 | 200 ± 310 | 27 ± 46 |
| | Mean Surface Area ± SD Per Object (µm²) | 420 ± 1.7E3 | 61 ± 150 | 4.5 ± 7.7 | 280 ± 350 | 3.4E3 ± 5.8E3 |
| *jrc_hela-3* | Total Volume (µm³) | 110 | 13 | 99 | 600 | 130 |
| | Total Surface Area (µm²) | 8.3E3 | 970 | 1.7E3 | 930 | 1.7E4 |
| | Number of Objects | 157 | 9 | 297 | 2 | 9 |
| | Mean Volume ± SD Per Object (µm³) | 0.68 ± 7.4 | 1.4 ± 3.8 | 0.33 ± 0.6 | 300 ± 400 | 15 ± 44 |
| | Mean Surface Area ± SD Per Object (µm²) | 53 ± 560 | 110 ± 290 | 5.6 ± 9.3 | 470 ± 530 | 1.8E3 ± 5.5E3 |
| *jrc_jurkat* | Total Volume (µm³) | 280 | 28 | 250 | 2.2E3 | 130 |
| | Total Surface Area (µm²) | 2.1E4 | 2.4E3 | 3.2E3 | 4.1E3 | 1.7E4 |
| | Number of Objects | 123 | 65 | 697 | 11 | 28 |
| | Mean Volume ± SD Per Object (µm³) | 2.3 ± 18 | 0.43 ± 1.1 | 0.36 ± 0.42 | 200 ± 100 | 4.8 ± 20 |
| | Mean Surface Area ± SD Per Object (µm²) | 170 ± 1.3E3 | 37 ± 94 | 4.6 ± 4.8 | 380 ± 220 | 600 ± 2.5E3 |
| *jrc_macrophage-2* | Total Volume (µm³) | 260 | 16 | 71 | 470 | 110 |
| | Total Surface Area (µm²) | 1.8E4 | 1.6E3 | 1.5E3 | 840 | 1.4E4 |
| | Number of Objects | 32 | 52 | 629 | 2 | 17 |
| | Mean Volume ± SD Per Object (µm³) | 8.2 ± 44 | 0.31 ± 1.8 | 0.11 ± 0.12 | 230 ± 330 | 6.4 ± 25 |
| | Mean Surface Area ± SD Per Object (µm²) | 550 ± 2.9E3 | 31 ± 180 | 2.5 ± 2.2 | 420 ± 530 | 840 ± 3.3E3 |
| *jrc-hela-1* | Total Volume (µm³) | 450 | | 360 | 1.9E3 | 320 |
| | Total Surface Area (µm²) | 3.4E4 | | 5.2E3 | 3.6E3 | 3.5E4 |
| | Number of Objects | 157 | | 799 | 9 | 180 |
| | Mean Volume ± SD Per Object (µm³) | 2.9 ± 17 | | 0.45 ± 1.5 | 210 ± 300 | 1.8 ± 23 |
| | Mean Surface Area ± SD Per Object (µm²) | 220 ± 1.3E3 | | 6.5 ± 1.3E3 | 400 ± 530 | 200 ± 2.5E3 |
| *jrc_choroid-plexus-2* | Total Volume (µm³) | 1.3E3 | | 1.0E3 | 2.9E3 | 180 |
| | Total Surface Area (µm²) | 1.0E5 | | 1.9E4 | 1.1E4 | 2.5E4 |
| | Number of Objects | 201 | | 2.5E3 | 36 | 120 |
| | Mean Volume ± SD Per Object (µm³) | 6.3 ± 89 | | 0.41 ± 2.9 | 81 ± 82 | 1.5 ± 16 |
| | Mean Surface Area ± SD Per Object (µm²) | 520 ± 7.3E3 | | 7.9 ± 55 | 300 ± 290 | 210 ± 2.2E3 |
| *jrc_cos7-11* | Total Volume (µm³) | 280 | | 130 | | |
| | Total Surface Area (µm²) | 2.3E4 | | 6.4E3 | | |
| | Number of Objects | 7.3E4 | | 1.3E3 | | |
| | Mean Volume ± SD Per Object (µm³) | 3.8E-3 ± 0.53 | | 0.11 ± 0.53 | | |
| | Mean Surface Area ± SD Per Object (µm²) | 0.31 ± 40 | | 5 ± 14 | | |

#### Supplementary Table 5- Select organelle measurements

Volume, instance count, and surface area measurements for select organelles in 7 different datasets. In many cases, the mean is small compared to the SD due to the wide range of object sizes for a small number of objects (i.e. ER and PM). For a complete list of measurements please refer to our data portal: openorganelle.janelia.org.

https://docs.aws.amazon.com/cli/latest/userguide/cli-chap-welcome.html.

57. Chamier, L. von, von Chamier, L., Laine, R. F. & Henriques, R. Artificial Intelligence for Microscopy: What You Should Know. doi:10.20944/preprints201902.0004.v2.
58. ISBI 2013 challenge: 3D segmentation of neurites in EM images. http://brainiac2.mit.edu/SNEMI3D/.
59. SegEM 3D EM segmentation challenge. https://segem.rzg.mpg.de//challenge/.
60. MICCAI Challenge on Circuit Reconstruction from Electron Microscopy Images. https://cremi.org/.
61. Haberl, M. G. *et al.* CDeep3M-Plug-and-Play cloud-based deep learning for image segmentation. *Nat. Methods* **15**, 677–680 (2018).
62. Bermudez-Chacon, R., Marquez-Neila, P., Salzmann, M. & Fua, P. A domain-adaptive two-stream U-Net for electron microscopy image segmentation. *2018 IEEE 15th International Symposium on Biomedical Imaging (ISBI 2018)* (2018) doi:10.1109/isbi.2018.8363602.
63. Roels, J., Hennies, J., Saeys, Y., Philips, W. & Kreshuk, A. Domain Adaptive Segmentation In Volume Electron Microscopy Imaging. *2019 IEEE 16th International Symposium on Biomedical Imaging (ISBI 2019)* (2019) doi:10.1109/isbi.2019.8759383.
64. ariadne.ai — AI-powered biomedical image analysis. https://ariadne.ai/.
65. Microscopy Image Analysis Software - Imaris - Oxford Instruments. https://imaris.oxinst.com/.
66. Object Research Systems (ORS) Inc, Montreal, Canada. *Dragonfly*. (2018).
67. Berg, S. *et al.* ilastik: interactive machine learning for (bio)image analysis. *Nat. Methods* **16**, 1226–1232 (2019).
68. Belevich, I., Joensuu, M., Kumar, D., Vihinen, H. & Jokitalo, E. Microscopy Image Browser: A Platform for Segmentation and Analysis of Multidimensional Datasets. *PLoS Biol.* **14**, e1002340 (2016).
69. Müller, A. *et al.* Three-dimensional FIB-SEM reconstruction of microtubule-organelle

interaction in whole primary mouse beta cells. 2020.10.07.329268 (2020) doi:10.1101/2020.10.07.329268.

70. Abadi, M. *et al.* TensorFlow: Large-Scale Machine Learning on Heterogeneous Distributed Systems. (2016).
71. Funke, J. Gunpowder. *GitHub* https://github.com/funkey/gunpowder.
72. Rocklin, M. Dask: Parallel Computation with Blocked algorithms and Task Scheduling. in (2015). doi:10.25080/majora-7b98e3ed-013.
73. Ledbetter, M. C. & Porter, K. R. A ‘MICROTUBULE’ IN PLANT CELL FINE STRUCTURE. *J. Cell Biol.* **19**, 239–250 (1963).
74. Kollmannsberger, P. *et al.* The small world of osteocytes: connectomics of the lacuno-canalicular network in bone. *New Journal of Physics* vol. 19 073019 (2017).
75. Bresenham, J. E. Algorithm for computer control of a digital plotter. *Seminal graphics* 1–6 (1998) doi:10.1145/280811.280913.
76. Zaharia, M. *et al.* Apache Spark: a unified engine for big data processing. *Commun. ACM* **59**, (2016).
77. Bogovic, J. A. *et al.* An unbiased template of the Drosophila brain and ventral nerve cord. 376384 (2018) doi:10.1101/376384.
78. Deng, J. *et al.* ImageNet: A large-scale hierarchical image database. in (2009). doi:10.1109/cvprw.2009.5206848.
79. Lehtinen, J. *et al.* Noise2Noise: Learning Image Restoration without Clean Data. in *International Conference on Machine Learning* 2965–2974 (PMLR, 2018).
80. Krull, A., Buchholz, T.-O. & Jug, F. Noise2Void - Learning Denoising From Single Noisy Images. in *2019 IEEE/CVF Conference on Computer Vision and Pattern Recognition (CVPR)* 2124–2132 (IEEE, 2020).
